# Modified defence peptides from horseshoe crab target and kill bacteria inside host cells

**DOI:** 10.1101/2021.06.27.450110

**Authors:** Anna S. Amiss, Jessica B. von Pein, Jessica R. Webb, Nicholas D. Condon, Peta J. Harvey, Minh-Duy Phan, Mark A. Schembri, Bart J. Currie, Matthew J. Sweet, David J. Craik, Ronan Kapetanovic, Sónia Troeira Henriques, Nicole Lawrence

**Affiliations:** Institute for Molecular Bioscience, Australian Research Council Centre of Excellence for Innovations in Peptide and Protein Science, The University of Queensland, Brisbane, Queensland, 4072, Australia; Institute for Molecular Bioscience, IMB Centre for Inflammation and Disease Research and Australian Infectious Diseases Research Centre, The University of Queensland, Brisbane, Queensland, 4072, Australia; Global and Tropical Health Division, Menzies School of Health Research, Darwin, Northern Territory, 0811, Australia; Australian Cancer Research Foundation / Institute for Molecular Bioscience Cancer Biology Imaging Facility, The University of Queensland, Brisbane, Queensland, 4072, Australia; School of Chemistry and Molecular Biosciences and Australian Infectious Diseases Research Centre, The University of Queensland, Queensland, Australia; Department of Infectious Diseases and Northern Territory Medical Program, Royal Darwin Hospital, Darwin, Northern Territory, 0811, Australia; Friedrich Miescher Institute for Biomedical Research, Basel, BS 4058, Switzerland; Queensland University of Technology, School of Biomedical Sciences, Translational Research Institute, Brisbane, Queensland, 4102, Australia

**Keywords:** Peptides, host defence, antimicrobial, macrophages, intracellular niche, uropathogenic *Escherichia coli*

## Abstract

Bacteria that occupy an intracellular niche can evade extracellular host immune responses and antimicrobial molecules. In addition to classic intracellular pathogens, other bacteria including uropathogenic *Escherichia coli* (UPEC) can adopt both extracellular and intracellular lifestyles. UPEC intracellular survival and replication complicates treatment, as many therapeutic molecules do not effectively reach all components of the infection cycle. In this study, we explored cell penetrating antimicrobial peptides from distinct structural classes as alternative molecules for targeting bacteria. We identified two β-hairpin peptides from the horseshoe crab, tachyplesin I and polyphemusin I, with broad antimicrobial activity toward a panel of pathogenic and non-pathogenic bacteria in planktonic form. Peptide analogues [I11A]tachyplesin I and [I11S]tachyplesin I maintained activity toward bacteria, but were less toxic to mammalian cells than native tachyplesin I. This important increase in therapeutic window allowed treatment with higher concentrations of [I11A]tachyplesin I and [I11S]tachyplesin I, to significantly reduce intramacrophage survival of UPEC in an *in vitro* infection model. Mechanistic studies using bacterial cells, model membranes and cell membrane extracts, suggest that tachyplesin I and polyphemusin I peptides kill UPEC by selectively binding and disrupting bacterial cell membranes. Moreover, treatment of UPEC with sublethal peptide concentrations increased zinc toxicity and enhanced innate macrophage antimicrobial pathways. In summary, our combined data show that cell penetrating peptides are attractive alternatives to traditional small molecule antibiotics for treating UPEC infection, and that optimization of native peptide sequences can deliver effective antimicrobials for targeting bacteria in extracellular and intracellular environments.

## INTRODUCTION

Antimicrobial resistance and tolerance pose a serious threat to global health, with the Centre for Disease Control and Prevention reporting more than 2.8 million cases of antibiotic-resistant infection and more than 35,000 associated deaths in the United States annually [1]. Due to intrinsic or acquired mechanisms of resistance, common and important antibiotics are becoming less effective [2, 3], leading to a growing need for new classes of antimicrobial compounds with different mechanisms of action to treat bacterial infections [4–7].

Antibiotic tolerance is a phenomenon where bacteria are genetically susceptible but phenotypically tolerant to antibiotic treatments [8, 9]. One mechanism of antibiotic tolerance is adoption of an intracellular lifestyle [10–12], where bacteria are protected from extracellular host defence mechanisms and antimicrobial therapy [11]. Pathogens with the ability to occupy intracellular niches promote their uptake by host cells, can escape or resist cellular antimicrobial mechanisms, adapt to the new host cell environment, and modulate host cell biology [10–12]. Depending on the bacterial pathogen, this intracellular niche can either be in the cytosol or inside vacuoles where they are shielded from antibiotic treatment and can facilitate recurrent infection (Fig. 1) [13, 14].

**Fig. 1.**
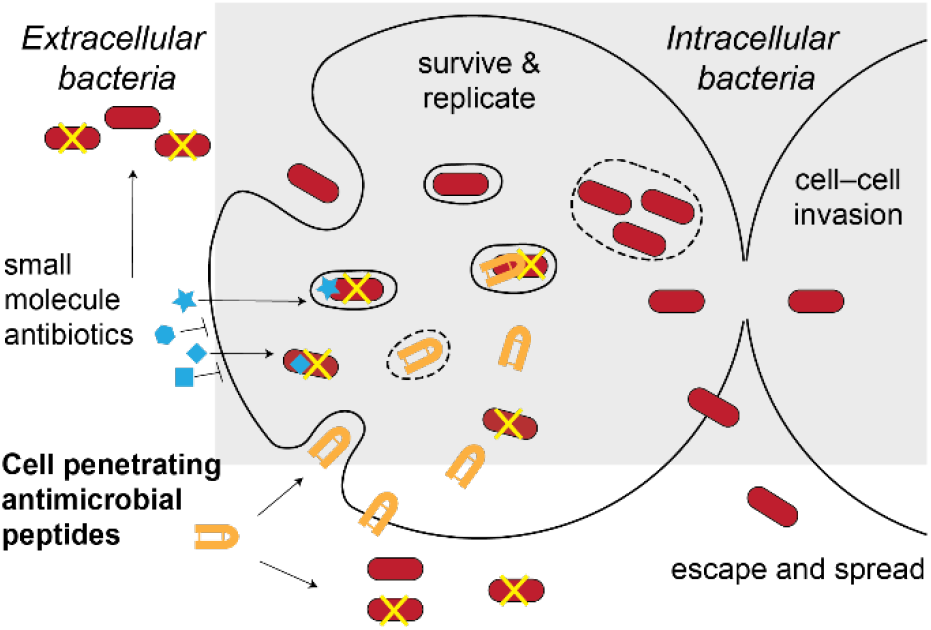
Challenges of targeting bacteria with an intracellular lifestyle. Pathogenic bacteria that survive and replicate inside cells can hide from cell-impermeable antimicrobial agents. Peptides able to cross host cell membranes to reach bacteria in the cytosol and/or vacuoles can overcome these challenges, representing candidates for developing new therapies for this challenging subset of bacteria.

The first barrier for a drug targeting intracellular bacteria is crossing the host cell membrane at non-toxic concentrations [10, 11, 15]. As host cell membranes are impermeable to most polar or charged molecules [11], this barrier precludes a large number of conventional therapeutics [10, 11]. The second barrier is the challenge of accessing bacteria located in the host cell cytosol, or sequestered inside membrane-bound vesicles, and maintaining antimicrobial activity in these environments (Fig. 1) [11, 15, 16]. Also, therapeutic molecules which can successfully reach the bacteria may not be at concentrations high enough to eradicate infection [11].

Classic examples of intracellular bacterial pathogens include *Mycobacterium tuberculosis*, *Salmonella enterica* serovar Typhimurium and *Burkholderia pseudomallei*. However, there is growing evidence that other bacteria, including *Pseudomonas aeruginosa, Staphylococcus aureus* and *Escherichia coli*, can occupy intracellular niches [17–20]. A pertinent example is uropathogenic *E. coli* (UPEC), which is the leading cause of urinary tract infections (UTIs) [21] that affect 40 – 50% of women during their lifetime [22, 23]. UPEC is primarily an extracellular pathogen that infects the bladder epithelium, but can also survive in intracellular niches such as within bladder epithelial cells where it proliferates to form intracellular bacterial communities [24, 25] and within neutrophils and macrophages, which are recruited to the bladder in response to UPEC infection [26–29]. The establishment of infection within immune cells is problematic as these cells can act as ‘Trojan horses’, leading to dissemination of the pathogen in the whole organism [10, 30]. Intracellular UPEC can also become dormant and form quiescent intracellular bacterial reservoirs which persist within the host cell, possibly contributing to recurrent UTIs [31]. Despite their previous use in the treatment of UTIs, widespread emergence of resistance to antibiotics such as trimethoprim-sulfamethoxazole, fluoroquinolones, and β-lactams has led to a decrease in their therapeutic efficacy [32–34].

Peptide-based therapeutics are an attractive alternative to conventional small molecule drugs. Current glycopeptide [35] and lipopeptide drugs [36] have potent antimicrobial activity, but are restricted to use as last-line therapies due to their narrow therapeutic index and undesirable off-target toxicity [37, 38]. For new molecule discovery, naturally occurring host defence peptides, collectively termed antimicrobial peptides, are gaining traction due to their broad spectrum of antimicrobial activity and efficacy at low and sub-micromolar concentrations. A subset of cell-penetrating antimicrobial peptides (herein referred to as CPPs) are especially attractive due to their ability to cross mammalian cell membranes and access intracellular targets [39–41].

CPPs are typically composed of 5–30 amino acids and have an amphipathic arrangement of hydrophobic and positively charged residues that promotes interaction with cell membranes [40, 42]. Furthermore, the arrangement of positively charged residues on one side of the peptide promotes electrostatic interactions with negatively charged phospholipids that are more commonly found in bacterial cell membranes, compared to neutral mammalian host cell membranes [40, 43–45]. Peptides are also thought to be less prone to common mechanisms of resistance observed for small molecule antibiotics because of their direct actions on microbial membranes, in contrast to affecting single, smaller molecular targets [39, 46].

Another strategy for avoiding antibiotic resistance is to enhance the effect of innate antimicrobial mechanisms. For example, innate immune cells such as macrophages and neutrophils use zinc poisoning as an antimicrobial weapon [47], and the zinc ionophore PBT2 reversed antibiotic resistance for several bacterial pathogens [48]. Interestingly, macrophage-mediated zinc toxicity is enhanced against *E. coli* mutants that are defective in zinc export (*ΔzntA*, *ΔzntR*), and also those with reduced membrane integrity (*ΔcpxR*, *ΔpstB*) [49]. Some pathogens, including *Salmonella* [50] and UPEC [49], can subvert intracellular zinc toxicity. We aimed to determine if amphipathic host defence peptides could enhance antimicrobial zinc toxicity due to their ability to reduce the membrane integrity of bacterial pathogens.

In this study, we examine the antimicrobial efficacy of host defence CPPs tachyplesin I and polyphemusin I toward pathogenic and non-pathogenic Gram-negative bacteria, and demonstrate enhanced zinc toxicity at sublethal peptide concentrations *in vitro*. Modification of native tachyplesin I improved selective toxicity toward UPEC, resulting in significantly reduced intracellular bacterial loads in an *in vitro* macrophage infection model. Our findings highlight the potential of CPPs as candidates for developing new therapeutics that target bacteria in both extracellular and intracellular environments.

## MATERIALS AND METHODS

### Bacterial strains

Pathogenic bacterial strains included in this study were: *B. pseudomallei* MSHR10517, MSHR2154 and MSHR1364 [51], *S.* Typhimurium SL1344, and UPEC ST131 strains EC958 (wild type, WT), EC958Δ*zntA* [49], and CTF073 [52]. Non-pathogenic strains were: *Burkholderia humptydooensis* (MSMB043) [51], and *E. coli* strains K-12 MG1655, ATCC 25922 and CSGC 7139 [53].

### Peptide synthesis and purification

Synthesis and purification of peptides were achieved using previously described methods [54]. Briefly, the peptides were synthesized using 9-fluroenylmethoxycarbonyl (Fmoc) solid-phase peptide synthesis on an automated peptide synthesizer (Symphony, Protein Technologies Inc, Tucson, USA). Rink amide resin was used for tachyplesin I, polyphemusin I, protegrin-I, Sub3, cyclic gomesin, while 2-chlorotrityl (2-CTC) resin was used for BP100 and analogues. The peptides were oxidized in 20% (*v/v*) solvent B with dropwise addition of a concentrated solution of iodine in acetic acid (persistent yellow colour) for 1 h. The reaction was quenched with ascorbic acid. Peptides were purified (>95%) using reverse-phase high performance liquid chromatography (Solvent A: H_2_O, 0.05% (*v/v*) trifluoracetic acid, solvent B: 90% (*v/v*) acetonitrile, 0. 05% (*v/v*) trifluoracetic acid). The correct peptide mass was confirmed using electrospray ionization mass spectrometry. The peptide concentration in aqueous solution was determined from the absorbance at 280 nm using the extinction co-efficient based on the contribution of tyrosine and tryptophan resides, as well as disulfide bonds [55].

### Peptide structure analysis

Correct fold and overall structure of peptides was determined by comparing one-dimensional ^1^H spectra with previously reported peptide structures. Additional two-dimensional structural information was obtained for [I11A]tachyplesin I and [I11S]tachyplesin I. Peptides (1 mg/mL) were dissolved in H_2_O/D_2_O (10:1, *v/v*) and the pH was adjusted to pH 4–5. One dimensional ^1^H spectra, and two-dimensional total correlated spectroscopy (TOCSY) and nuclear Overhauser effect spectroscopy (NOESY) were acquired at 298K with a Bruker Avance III HD 600 MHz NMR spectrometer. Spectra were referenced externally to 2,2-dimethyl-2-silapentone-5-sulfonate (DSS) at 0 ppm. After processing with TOPSPIN 3.6 (Bruker), spectra were assigned using CcpNmr [56]. Secondary αH shifts were calculated as the difference between observed αH chemical shifts and those of the corresponding residues in random coil peptides [57].

### Antimicrobial activity screen

The minimum inhibitory concentrations (MICs) of the peptides against *Burkholderia, E. coli,* UPEC and *Salmonella* strains was determined using a plate-based broth microdilution method [58]. Overnight bacterial cultures were subcultured until mid-log phase then diluted to OD_600_ = 0.01 in broth (Mueller Hinton Broth for *Burkholderia* strains, or lysogeny broth (LB) for *E. coli*, UPEC and *S.* Typhimurium strains). Peptides were serially diluted starting at 64 µM. Plates were incubated at 37°C for 24 h and the MIC was determined by the lowest concentration of compound that inhibited visible bacterial growth. Data represent the MIC determined from three independent experiments for each bacterial strain.

### Bone marrow-derived macrophage cytotoxicity

Bone-marrow cells were harvested from the femurs and tibias of male C57Bl/6 mice aged between 7 and 12 weeks old, as previously described [59]. All studies involving animals were approved by The University of Queensland animal ethics committee (AEC IMB/123/18). Bone-marrow derived macrophages (BMMs) were obtained by *in vitro* differentiation of the bone-marrow cells in complete RPMI 1640 (media supplemented with 10% heat-inactivated foetal bovine serum, 2 mM _L_-glutamine, 50 U/ mL penicillin and 50 μg/mL streptomycin with recombinant human CSF-1 (10,000 U/mL, UQ Protein Expression Facility) [59]. BMMs were differentiated at 37 °C in a humidified incubator at 5% CO_2_ for six days prior to experimentation.

Day six BMMs were seeded at a density of 80,000 cells/well in a 96-well flat bottom cell culture plate. Cells were left overnight in complete RPMI to rest. On day seven, the media was removed, and cells were washed twice with phosphate buffered saline (PBS). Cell toxicity was determined following peptide treatment using a resazurin assay as before [60]. Serum-free and antibiotic-free media (RPMI supplemented with 2 mM L-glutamine and recombinant human CSF-1 (10,000 U/mL)) was added to the well with serial dilutions of compounds starting at 64 µM. Untreated cells and cell lysed by 0.1% (*v/v*) Triton X-100 were used as controls to establish 0% and 100% of cell death, respectively. The plate was returned to the incubator for 2 h, before addition of 0.01% (w/v) resazurin and return to the incubator for a further 22 h. Fluorescence emission intensity (excitation at 560 nm, emission at 585 nm) was then measured on a Tecan Infinite M1000 Pro. The percentage of cell death was calculated compared to 100% death induced by the Triton X-100 control. Data were collected from three independent experiments.

### Red blood cell hemolysis assay

Red blood cells (RBCs) were isolated from healthy adult donors and washed three times in PBS (centrifuged at 500 *g* for 1 min). Peptides were diluted in PBS and incubated with 0.25% (*v/v*) RBCs at 37 °C for 1h. Samples with 0.1% (*v/v*) Triton X-100 were included as a 100% hemolysis control, and PBS was included as a 0% hemolysis control. Plates were centrifuged (500 *g* for 5 min) to pellet the RBCs. The supernatant was transferred to a flat-bottomed 96-well plate, and the absorbance was read at 415 nm on a Tecan Infinite M1000 Pro to detect released hemoglobin. The percentage hemolysis was calculated based on the hemolysis of the 0.1% Triton X-100 sample [61]. Data were collected from a single experiment using RBCs from two individual donors.

### Synthetic lipid vesicle preparation

The synthetic lipids 2-oleoyl-1-palmitoyl-sn-glycero-3-phosphocholine (POPC), POPC/ 1-palmitoyl-2-oleoyl-sn-glycero-3-phosphoethanolamine (POPE) (4:1 molar ratio mixture) and POPC/ 1-palmitoyl-2-oleoyl-sn-glycero-3-phosphoglycerol (POPG) (4:1 molar ratio mixture) were prepared as a lipid film by mixing in chloroform and then drying under both nitrogen gas and a vacuum desiccator overnight. The lipid film was subjected to repeated freeze/ thaw cycles then extruded at 1 mM in surface plasmon resonance (SPR) running buffer (10 mM HEPES, 150 mM NaCl, pH 7.4) through membranes with 50 nm pores to form small unilamellar lipid vesicles as previously described [62].

### Membrane extract vesicle preparation

To extract bacterial cell membranes, mid-log phase bacterial cultures were centrifuged at 2000 *g* for 5 min. The supernatant was removed, and cells were washed in PBS. Bacterial cells were then centrifuged at 2000 *g* for 5 min and the supernatant was removed. The membranes of RBCs and BMMs were extracted by similar methods. After washing with PBS and removing the supernatant, 110 µL of methanol and 390 µL of methyl tert-butyl ether (MTBE) with 0.01% butylated hydroxytoluene (BHT) was added to each sample [63]. Samples were vortexed to mix, followed by shaking incubation for 1 h. Next, 100 µL of 150 mM of ammonium acetate was added, and the samples were vortexed for 20 s before 5 min of centrifugation at 2000 *g*. The top (MTBE) phase was removed and stored in glass vial, taking care not to disturb the aqueous phases. Cell membrane extracts were dried under nitrogen gas and a vacuum desiccator overnight. Vesicles were prepared using the same method of freeze/thaw and extrusion as described for the synthetic lipid vesicles.

### Surface plasmon resonance

SPR was used to measure the affinity of peptides for lipid membranes (Biacore T200). SUVs prepared as described above were deposited onto a Biacore L1 chip at a flow rate of 2 µL/min for 40 min, to reach a steady-state plateau and surface coverage with lipid bilayer. Serial dilutions of peptides (in SPR running buffer) were injected and diffused over the lipid bilayer at 5 µL/ min for 180 s. The peptide–lipid disassociation was monitored for 600 s. Following the association phase, the response units (RU) were converted to peptide to lipid ratio (P/L, mol/mol) for each peptide dilution, assuming 1 RU = 1 pg mm^-2^. The maximum P/L ratio (P/L max) was determined by fitting P/L dose response curves (saturation binding with Hill slope) [64].

### Bacterial cell permeability

SYTOX™ Green nucleic acid stain in combination with flow cytometry was used to investigate the integrity of bacterial outer and inner membranes [65]. Samples were prepared a 96–well plate, performing serial dilutions from the determined MIC. Peptide vehicle treated (10% water) bacteria were used as the 0% permeability control and bacteria treated with 50% isopropanol were used as the 100% permeabilized control.

After 1 h of treatment, 200 µL of ice-cold PBS was added to each well. Samples were transferred to FACS tubes and kept at 4 °C while previous samples were run. 1.3 µM final concentration of SYTOX™ Green was added to each sample prior to loading into a CytoFLEX S Flow Cytometer (Beckman Coulter). 100,000 gated events were recorded for each condition. The percentage of permeabilized cells was calculated relative to the isopropanol control. Data were collected from three independent experiments.

### Flow cytometry to measure peptide internalization

Peptides were labelled with Alexa Fluor® 488 (A488) 5-sulfodichlorophenol ester (SDP) (Thermo Fisher Scientific) using previously described methods [66]. Peptide (0.5–1 mg) was dissolved in dimethylformamide, A488 5-SDP (dissolved to 10 mM in dimethylsulfoxide (DMSO)) and N,N-diisopropylethylamine at a ratio of 78:20:2 (*v/v/v*). The peptide labelling reaction was performed at room temperature, protected from light for 2 h. A488 labelled peptide was purified using analytical scale HPLC. The correct peptide mass was confirmed using mass spectrometry as above. Stocks of labelled peptide were prepared, and their concentration quantified by measuring the absorbance at 495 nm (ε_495_ = 73,000 M^−1^ cm^−1^).

Fluorescently labelled peptide was used to determine the percentage of BMM cells with peptide internalized. BMMs were seeded and incubated as per the cytotoxicity assay. Media was removed and replaced with 90 µL serum free media and 10 µL of labelled peptide stock. Cells were incubated at 37°C and 5% CO_2_ for 1 h. Cells were lifted from the plate using ice cold PBS, transferred to tubes, and analysed using a CytoFLEX S Flow Cytometer. Trypan blue (0.02% w/v final) was added to each sample to quench the fluorescence of labelled peptide accessible to aqueous environment (e.g., bound onto the surface of cells, but not internalized) before the recording was repeated [67]. The percentage of fluorescent cells with fluorescence above a background threshold for cells without peptide was determined from 10,000 gated events. Data were collected from three independent experiments.

### Live cell imaging to examine internalization of tachyplesin into BMMs

Day 7 BMMs were seeded at 80,000 cells per well in a Nunc™ Lab-Tek™ Chambered coverglass and incubated overnight at 37 °C and 5% CO_2_. Before the assay, cells were washed with PBS, then incubated with serum free RPMI media containing 2 µM tachyplesin I labeled with A488 and 100 µg/mL dextran-tetramethylrhodamine (TMRE) (Thermo Fisher Scientific) for 15 min. The media was removed and replaced with FluoroBrite DMEM media (Thermo Fisher Scientific). Cells were incubated at 37 °C and 5% CO_2_ during imaging. Localization of A488 labelled tachyplesin I inside BMMs was examined using a Zeiss LSM 880 inverted confocal microscope using a 63x 1.4 NA C-Plan Apochromat oil immersion objective and acquired with Zen Black imaging software with a pinhole of 1 AU set for all channels and a pixel dwell time of 0.25 µs. A488 labelled peptide was excited with the 488 nm line of an Argon Ion laser and the fluorescence emission detected with the internal GaAsP detector gated to 492–556 nm, whereas dextran-TMRE was excited with the 561 nm line of a solid state laser and fluorescence emission detected with the internal GaAsP detector gated to 566–685 nm with a gain voltage of 500. Images were post processed using Fiji software [68].

### Microscopy to determine location of peptides inside infected BMMs

Samples for microscopy were prepared using day 7 BMMs seeded at 80,000 cells per well in a Nunc™ Lab-Tek™ Chambered coverglass. Bacterial infection assays were performed as previously described [26]. Briefly, EC958 was cultured overnight at 37 °C in LB under static growth conditions to enrich for type 1 fimbriae expression, which was confirmed by yeast-agglutination as previously described [69]. Cells were regrown for 4 h under the same conditions and then used to infect BMMs (MOI 1:100) for 1 h. Media was removed and replaced with fresh media containing 200 µg/mL gentamicin for an additional 1 h. The media was then removed and replaced with media containing 20 µg/mL gentamicin and 8 µM of fluorescently A488-labelled [I11A]tachyplesin I or [I11S]tachyplesin I. BMMs were treated with peptide for 1 h before being washed three times with PBS. Cells were fixed used 4% paraformaldehyde (15 min at room temperature). The fixative was removed and replaced with PBS. Cells were stained with Wheat Germ Agglutinin (WGA)-TMRE (Thermo Fisher Scientific) and 4′,6-diamidino-2-phenylindole (DAPI) (Thermo Fisher Scientific).

Multichannel confocal images were acquired using a Zeiss LSM 880 inverted confocal microscope using a 63x 1.4NA C-Plan Apochromat oil immersion objective and acquired with Zen Black imaging software with a pinhole of 1 AU set for all channels and a pixel dwell time of 0.5 μs. Peptide (A488 labeled) was excited with the 488 nm line of an Argon Ion laser and its fluorescence emission detected with the internal GaAsP detector gated to 492-623 nm, while WGA-TMRE and DAPI were co-excited with the 405 nm and 561 nm lines of individual lasers and detected with the internal GaAsP detectors gated to 410–483 nm, and 566–685 nm with voltage gains of 650 and 713 respectively. Images were acquired as Z-stacks covering the full height of the cell at intervals of 250 nm and are displayed as maximum intensity projections.

### Co-treatment of planktonic EC958 with peptides and metal ions

A checkerboard strategy was employed to examine the effect of peptide co-treatment with metal ions on the growth of planktonic bacteria. Peptides were serially diluted with zinc sulfate, copper (II) sulfate or iron (II) sulfate on a 96-well plate. Starting concentrations of peptides against each bacterial strain was determined from the antimicrobial peptide screen. The broth microdilution assay was performed as described above. After the addition of bacteria, the plate was placed in a POLARstar Omega (BMG LABTECH) at 37 °C. Absorbance was read at 600 nm every 30 min for 12 h, with orbital shaking performed before the readings. Data points were collected from three independent experiments for each bacterial strain.

### Peptide treatment of bone marrow-derived macrophages infected with EC958

Day 7 BMMs were infected with static cultures of type I fimbriae enriched EC958 WT or EC958Δ*zntA* bacteria (enriched as described for EC958 WT, with MOI 1:100) for 1 h. Media (RPMI supplemented with 10% FCS, 2 mM Glutamax and colony stimulating factor 1 (CSF-1) was replaced with fresh media containing 200 µg/mL gentamicin for an additional 1 h. The media was then removed and replaced with media containing 20 µg/mL gentamicin and tachyplesin I, [I11A]tachyplesin I or [I11S]tachyplesin I. At 6 h post infection (4 h post peptide treatment) samples of each condition were lysed using 0.1% (*v/v*) Triton X-100 in PBS. Dilutions of the lysates were plated on agar (LB agar for EC958 WT cells and LB agar with 30 mg/chloramphenicol for EC958Δ*zntA)*. This step was repeated 12 h post infection. Agar plates were placed in a 37°C incubator overnight (∼18 h). CFU were determined by enumerating the number of colonies per condition. Data were collected from three independent experiments.

For each experiment, a lactate dehydrogenase activity assay (LDH) (using the CytoTox 96® Non-Radioactive Cytotoxicity Assay LDH Kit (Promega)) was performed in parallel to determine host cell toxicity during the infection. BMMs were infected and treated in the same manner as the infection plates. Controls for 100% lysis were included by adding 0.1% (*v/v*) Triton X-100 into control wells and incubating for 10 min before removing the supernatant sample. At 6 and 12 h post infection, 25 µL of the supernatant from each condition was removed and transferred to a new plate. The LDH assay was then performed as per the manufacturer’s instructions. Absorbance at 480 nm was read on a Tecan infinite M1000Pro and the percentage cell death was calculated using the ratio of media subtracted conditions divided by subtracted absorbance of 100% lysis controls.

## RESULTS AND DISCUSSION

### Host defence CPPs

The panel of peptides in this study (see Fig. 2) comprises host defence CPPs with reported membrane-active mechanisms of action [60, 65, 70–72]. It includes peptides representing the two major secondary structures (i.e., α-helix and β-hairpin), with differences in amino acid sequence, amount of positive charge, and distinct amphiphilicity (hydrophobic moment). These properties are likely to impact antimicrobial activity [73]. Two analogues were prepared for tachyplesin I by modifying an Ile residue (position 11) to either Ala or Ser to produce [I11A]tachyplesin I and [I11S]tachyplesin I, respectively. Substitutions at this position have been previously shown to improve selectivity for bacterial membranes compared to mammalian cells [41, 71].

**Fig. 2.**
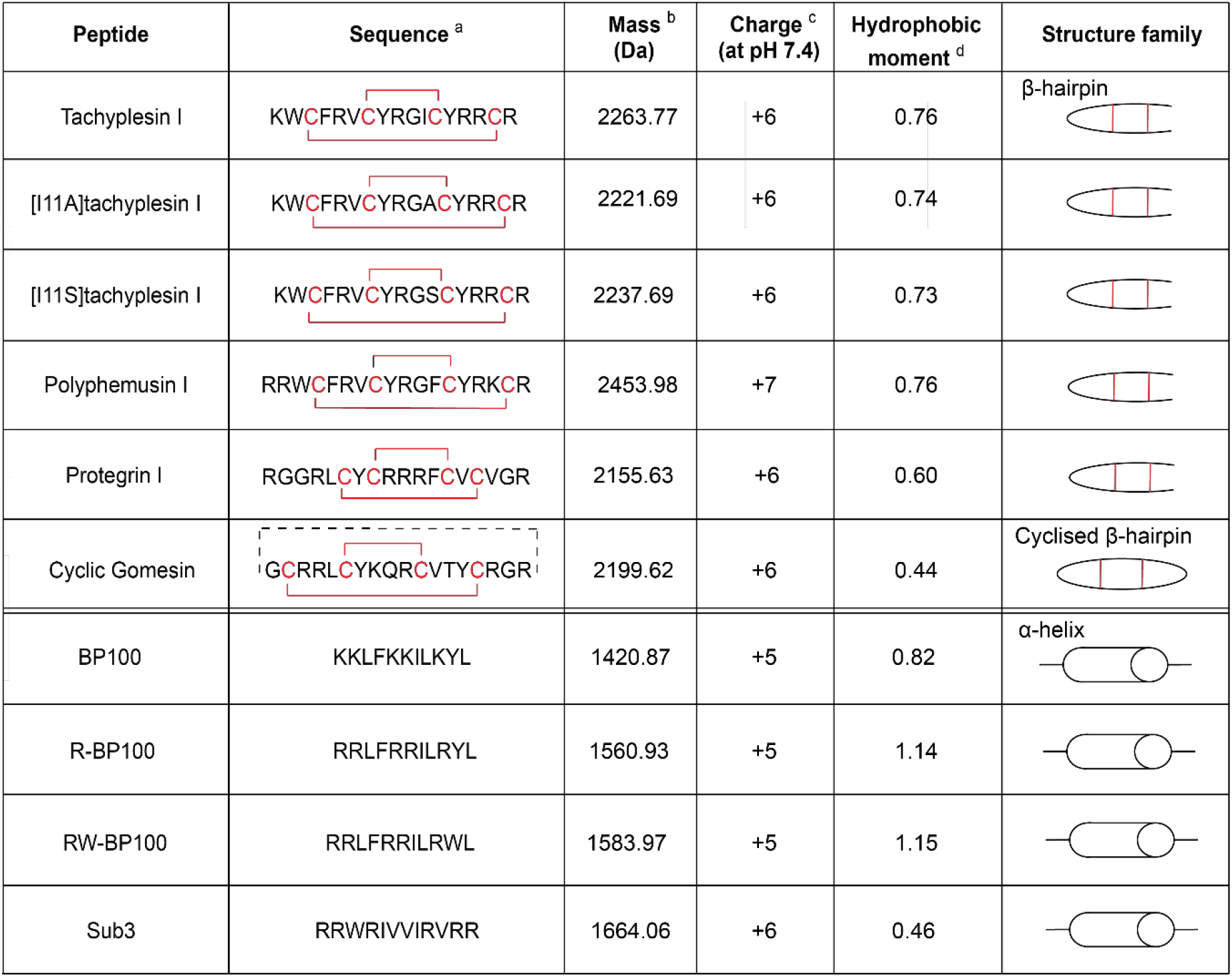
Characteristics of the peptide panel used for antimicrobial susceptibility screening. ^a^ Cys residues and disulfide bonds are shown in red. Backbone cyclisation is illustrated by a dashed black line. Sequence reference: tachyplesin I and [I11A]tachyplesin I [71], polyphemusin I and protegrin I [70], cyclic gomesin [65], BP100 and analogues [60], Sub3 [72]. ^b^ Peptide mass (Da) is the average theoretical mass calculated from the amino acid sequence. ^c^ Peptide overall charge was calculated from contributions of charged amino acid side chains. ^d^ Hydrophobic moment was calculated from the amino acid sequence using the hmoment function of the R package [74].

### Comparison of the overall secondary structure of tachyplesin I and analogues

Differences in peptide structure can affect peptide–membrane interactions and activity; therefore, structural studies were undertaken to investigate whether the Ile to Ser substitution affected the overall structure of [I11S]tachyplesin I compared to the native tachyplesin I. The secondary αH chemical shift for tachyplesin I [41, 71, 75] and [I11A]tachyplesin I [71] were previously described, and are compared here to the novel secondary αH chemical shift for [I11S]tachyplesin I (Fig. 3A).

**Fig. 3.**
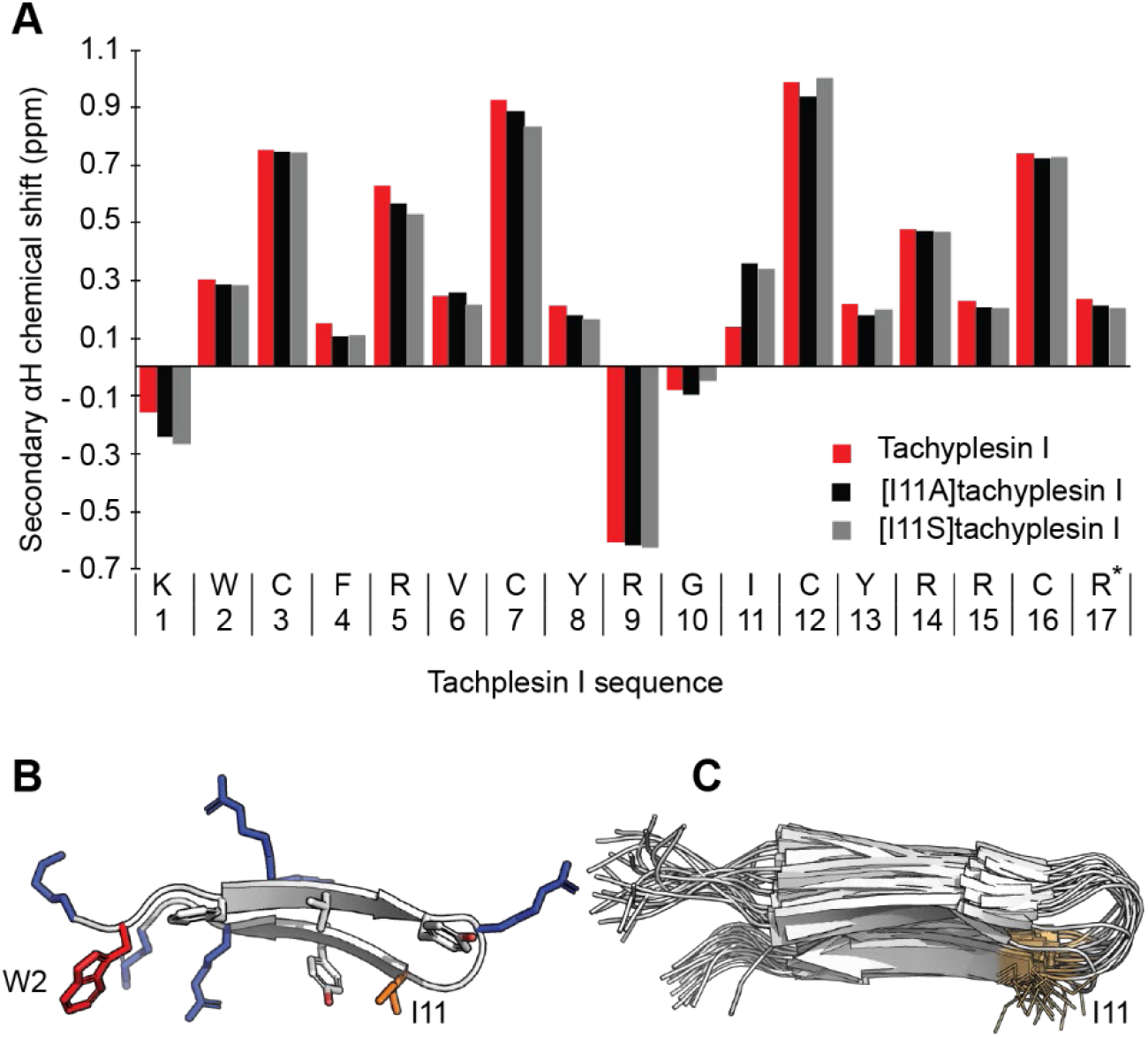
Structural characteristics of tachyplesin I and analogues. (A) Secondary αH chemical shift at 298 K determined from 1H NMR spectra. The secondary αH shifts of tachyplesin I (red), [I11A]tachyplesin I (black) and [I11S]tachyplesin I (grey) were calculated as the difference between observed αH chemical shifts and those of the corresponding residues in random coil peptides [72]. (B) Cartoon representation of tachyplesin I (PDB ID: 2RTV) showing amphipathic arrangement of positively charged (blue) and hydrophobic residues (orange). W2 and I11 are situated at opposite ends of the peptide. (C) Overlay of the 20 lowest energy forms of tachyplesin I (PDB ID: 2RTV) showing the consistent solvent exposed positioning of the side chain of I11.

All three peptides show a defined β-hairpin structure, with a high conservation of αH chemical shifts. [I11A]tachyplesin I and [I11S]tachyplesin I show similar deviation at position 11. In the native tachyplesin I peptide, the side chain of I11 is positioned at the opposite end of the peptide to the hydrophobic residue W2 that inserts into lipid bilayers [41] (Fig. 3B) and is solvent-exposed in the lowest energy structures (Fig. 3C). Ala has a shorter hydrophobic side chain than Ile, and Ser has a polar side chain with a hydroxyl group, which decreases local hydrophobicity and can establish H-bonds with phospholipid headgroups. Thus, while the overall structure of the peptide is conserved, the different characteristics of the amino acid side chain at position 11 (Ala or Ser) may affect peptide binding and insertion into biological membranes, possibly enhancing antibacterial activity.

### Peptide susceptibility screen against planktonic bacteria

The peptides were tested for activity against representative pathogenic and non-pathogenic bacteria. The minimum inhibitory concentration (MIC) was determined for planktonic bacteria (Table 1) and used to identify the most potent candidates for progression in this study. The pathogenic bacteria – *B. pseudomallei*, UPEC strain EC958, and *S.* Typhimurium (shaded in grey, Table 1) – share the ability to survive and replicate within host cells, making them the most suitable bacteria for this study. CFT073 was included to account for UPEC strain differences in virulence and innate immune responses [76], particularly with respect to interactions with macrophages [77, 78]. The non-pathogenic bacteria – *Burkholderia humptydooensis* and *E. coli* strains – (unshaded, Table 1) were included as control strains to illustrate the broader context of antimicrobial susceptibility and provide information on the specificity and mechanism of action. Notably, we have previously demonstrated that the non-pathogenic species *B. humptydooensis* recapitulates the antimicrobial susceptibility of *B. pseudomallei,* and can be used as a lower-risk group model organism in planktonic studies [51].

**Table 1:** MIC of peptides against pathogenic and non-pathogenic bacteria strains, compared to toxicity in host mammalian cells.

| Antimicrobial activity: MIC ($\mu\text{M}$)<sup>a</sup> | | | | | | | | | | Mammalian cell toxicity CC50 ($\mu\text{M}$) | |
| --- | --- | --- | --- | --- | --- | --- | --- | --- | --- | --- | --- |
| Peptide |  | *Burkholderia pseudomallei*<sup>b</sup> | *Burkholderia humptydooensis*<br>MSMB 043 | *Uropathogenic E. coli* |  | *Non-pathogenic E. coli* |  |  | *Salmonella* Typhimurium<br>SL1344 | BMM<sup>c</sup> | RBC<sup>d</sup> |
|  |  |  |  | EC958 | CFT073 | K-12<br>MG1655 | ATCC<br>25922 | CSGC<br>7139 |  |  |  |
| β-hairpin | Tachyplesin I | > 64 | 32 | 0.25 – 0.5 | 0.25 – 0.5 | 0.125 – 0.25 | 0.125 – 0.5 | 0.5 | 0.25 – 0.5 | 11.1 ± 1.3 | 42.1 ± 5.9 |
| | [I11A]tachyplesin I<sup>e</sup> | nd | nd | 0.5 – 2 | nd | 0.5 – 2 | nd | nd | nd | 26.7 ± 1.5 | >> 128 |
| | [I11S]tachyplesin I<sup>e</sup> | nd | nd | 0.5 – 2 | nd | 0.5 – 2 | nd | nd | nd | 29.3 ± 1.4 | >> 128 |
| | Polyphemusin-I | > 64 | 32 – 64 | 1 – 2 | 0.5 – 2 | 0.5 – 1 | 4 | 0.5 – 1 | 1 – 4 | 12.6 ± 2.3 | 52.3 ± 7.0 |
|  | Protegrin-I | > 64 | 64 | 4 – 16 | 16 – 64 | 2 – 4 | 4 – 8 | 2 – 4 | 8 – 64 | nd | nd |
|  | Cyclic gomesin | nd | > 64 | 2 – 16 | 4 – 8 | 8 | 2 | 1 – 2 | 32 |  |  |
| α-helix | BP100 | nd | > 64 | 16 – 32 | 32 – 64 | 16 – 64 | 1 – 2 | 0.5 – 1 | 8 – 16 | | |
|  | R-BP100 | nd | > 64 | 16 – 32 | 16 – 32 | 16 – 64 | 2 – 4 | 1 – 2 | 8 – 16 |  |  |
|  | RW-BP100 | nd | > 64 | 32 – 64 | 32 – 64 | 32 – > 64 | 2 – 16 | 1 – 4 | 8 – 32 |  |  |
|  | Sub3 | nd | > 64 | 16 – 32 | 32 – 64 | 8 – 16 | 2 – 4 | 0.5 – 1 | 32 – 64 |  |  |

Grey columns denote pathogenic bacteria, nd indicates where an MIC was not determined.
<sup>a</sup> MICs were determined using broth microdilution of planktonic bacteria in growth phase. Data represent the MIC range determined from a minimum of three independent experiments. 64 $\mu\text{M}$ was the highest concentration tested.
<sup>b</sup> MIC was determined for *B. pseudomallei* strains MSHR1364, MSHR2154 and MSHR10517.
<sup>c</sup> BMMs were treated with serial dilutions of tachyplesin I, polyphemusin I, [I11A]tachyplesin I or [I11S]tachyplesin I at 37 °C for 24 h and viability was determined using resazurin (0.01 % w/v). Triton X-100 (0.1% v/v) was included to establish 100% cell death. The cytotoxic concentration required to kill 50% of the cells (CC50) was determined from dose response curves, where the nonlinear regression was fit using saturation binding with Hill slope (GraphPad Prism 8). Data represent a minimum of three independent experiments.
<sup>d</sup> RBCs were treated with serial dilutions of tachyplesin I, polyphemusin I, [I11A]tachyplesin I or [I11S]tachyplesin I at 37 °C for 1 h. Triton X-100 (0.1% v/v) was included to establish 100% hemolysis. Hemolysis was determined by monitoring haemoglobin in the supernatant. The concentration required to lyse 50% of the cells (CC50) was determined from the dose response curves, where the nonlinear regression was fit using [inhibitor] vs response with a variable slope (GraphPad Prism 8). Data represent a single experiment with RBCs isolated from two healthy donors. 128 $\mu\text{M}$ was the highest concentration tested.
<sup>e</sup> [I11A] and [I11S]tachyplesin were designed as non-toxic analogues for targeting intracellular bacteria. Therefore, antimicrobial activity was determined for EC958 and control non-pathogenic MG1655 strains.

Tachyplesin I and polyphemusin I were the two most active peptides across the entire bacteria panel (Table 1). Antimicrobial activity of [I11A]tachyplesin I [71], and [I11S]tachyplesin I (new analogue) was tested against EC958 and non-pathogenic MG1655. Both analogues retained low micromolar potency toward these bacterial strains.

Overall, the β-hairpin peptides (tachyplesin I and analogues, polyphemusin I, protegrin I, and cyclic gomesin) were more active against the tested bacterial strains than the α-helical peptides (BP100 and analogues, and Sub3), suggesting that the β-hairpin structure contributes to more potent antimicrobial activity; especially toward pathogenic UPEC and *S.* Typhimurium strains. Comparing the β-hairpin peptides, there appeared to be a relationship between a higher hydrophobic moment value (a measure of amphiphilicity, where a large hydrophobic moment indicates a structure that is predominantly hydrophobic on one side and predominantly hydrophilic on the other) [79] and a lower MIC. For example, polyphemusin I, and tachyplesin I and analogues (hydrophobic moment range 0.73 – 0.76) were consistently more active than protegrin I and cyclic gomesin (hydrophobic moments of 0.60 and 0.44). All tested peptides had an overall positive charge (+5 to +7) and no distinction in antimicrobial activity was correlated with difference in charge within this range.

### Host cell toxicity

When developing new therapies to target bacteria within intracellular niches, it is important to consider toxicity of the active antimicrobials against host cells. To this end, we determined the toxicity of the most potent antimicrobial peptides from the planktonic activity assays (Table 1); tachyplesin I, polyphemusin I, [I11A]tachyplesin I and [I11S]tachyplesin I, toward human RBCs and murine BMMs.

The two tachyplesin I analogues, [I11A] and [I11S], were the least toxic peptides, with CC50 (peptide concentration required to kill 50% of BMMs) values approximately 2-fold higher than tachyplesin I and polyphemusin I. Notably, these peptide concentrations are 10- to 100-fold higher than the concentrations required to inhibit growth of 100% of *E. coli* and *S.* Typhimurium strains (see Table 1). For RBC toxicity, we confirmed previously reported concentrations of tachyplesin I [66, 71, 80] and polyphemusin I [80, 81] required to induce hemolysis, and reduced toxicity of [I11A]tachyplesin I [71] (see Table 1). Incubation with 128 µM [I11S]tachyplesin I and [I11A]tachyplesin I resulted in < 50% lysis of RBCs; therefore, the tachyplesin I analogues were at least 3-fold less hemolytic than the native tachyplesin I (42.1 ± 5.9 µM) and polyphemusin I (52.2 ± 7.0 µM). These differences in effective concentration underline the greater therapeutic possibilities of [I11A]tachyplesin I and [I11S]tachyplesin I during bacterial infection as they are better tolerated by mammalian cells.

### Peptide–lipid binding affinity

Peptide–lipid membrane binding studies were performed using SPR to determine whether differences in selective antimicrobial activity of tachyplesin I, polyphemusin I and the two tachyplesin I analogues relate to differences in their ability to interact with the phospholipid bilayer in cell membranes. Lipid model membranes composed of synthetic zwitterionic phospholipid POPC were used to represent the neutral outer leaflet in the cell membrane of healthy eukaryotic cells. Phospholipids containing phosphatidylethanolamine (PE)- or phosphatidylglycerol (PG)-headgroups are the major lipid components of bacterial membranes [43]; therefore, model membranes composed of POPC/POPE (4:1 molar ratio) and POPC/POPG (4:1 molar ratio) were also investigated.

All four peptides had the highest affinity for POPC/POPG (4:1), followed by POPC, then POPC/POPE (4:1) model membranes (Fig. 4), as shown by their sensorgrams obtained with peptide at 16 μM, dose-response curves, and quantified by peptide-to-lipid ratio obtained at binding saturation (P/L_max_). This trend suggests that affinity for phospholipids containing PG headgroups may drive peptide interaction with bacterial membranes. P/L_max_ values (Fig. 4B) show that tachyplesin I has the greatest binding affinity to all three model membranes; approximately 1.5-fold higher than polyphemusin I, [I11A]tachyplesin I and [I11S]tachyplesin I for POPC/POPG, and 2-fold higher for POPC. This is further demonstrated from the peptide–membrane binding characteristics, as shown by the representative sensorgrams for 16 µM peptides in Fig. 4C. The peptide–lipid dissociation constants (k_d_) were calculated by fitting the dissociation phase of the sensorgram obtained with peptide at 16 μM and were found to be similar across the peptides toward each of the model membranes (ranging from 0.005 – 0.009 s^-1^); this suggests that variations in the peptide–lipid binding affinity relate to differences in the association rate (e.g. tachyplesin I associates with lipid bilayers at a faster rate than the other peptides).

**Fig. 4.**
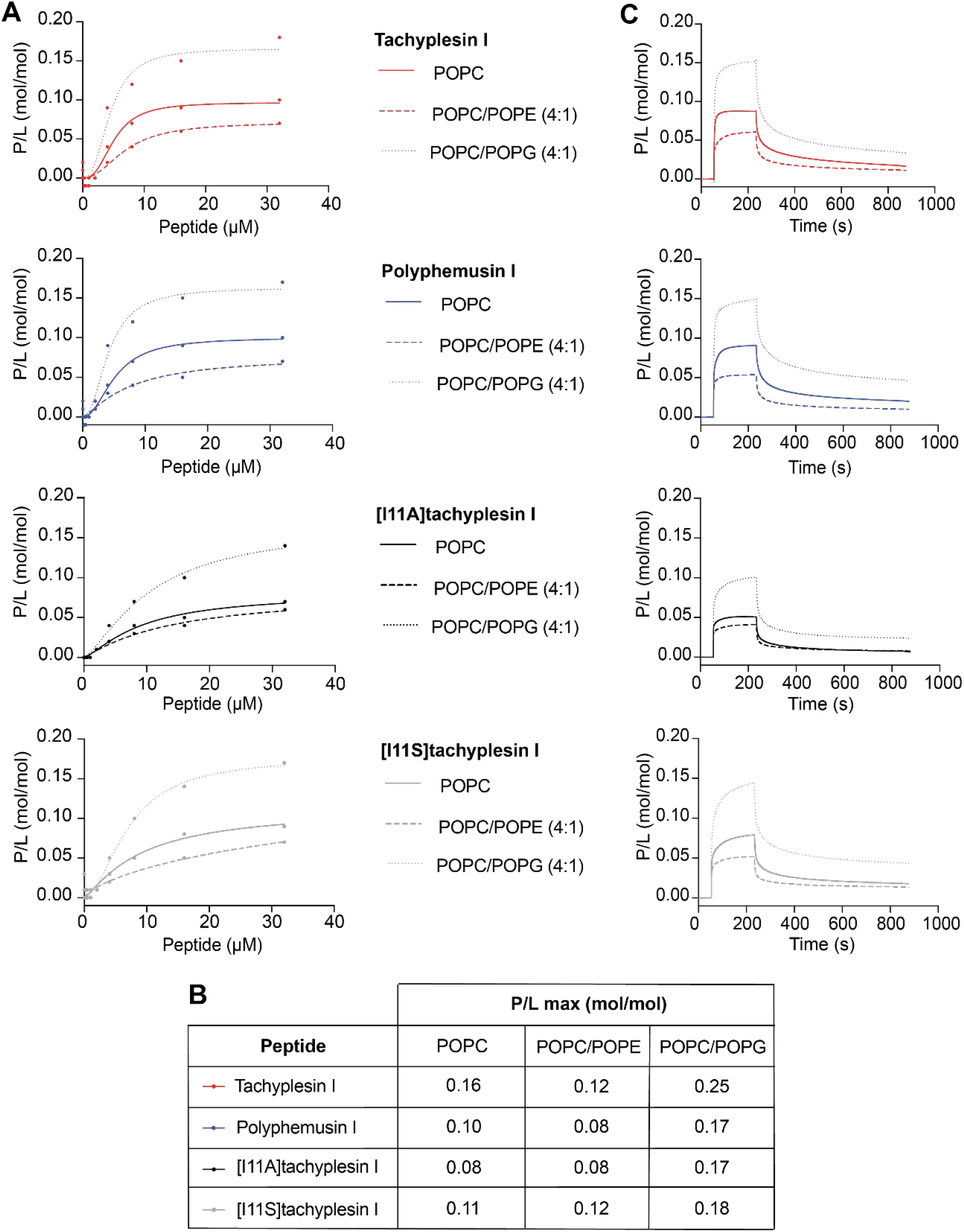
Membrane binding affinity and dose-response curves of tachyplesin I, polyphemusin I, [I11A]tachyplesin I and [I11S]tachyplesin I toward synthetic lipid model membranes. Comparison of the binding affinity of each peptide for model membranes composed of POPC, POPC/POPE (4:1) and POPC/POPG (4:1). SPR sensorgrams were obtained for peptides injected over lipid bilayers deposited on an L1 chip for 180 s, with dissociation monitored for 600 s. The response units at the end of the association phase were converted to peptide-to-lipid molar ratios (P/L (mol/mol)). (A) Dose response curves allow comparison of the amount of peptide bound to different lipid mixtures. (B) The maximum P/L ratio (P/L max) was determined by fitting P/L doses response curves (saturation binding with Hill slope, GraphPad Prism 8). (C) Representative sensorgrams for 16 µM peptide show similar peptide-membrane dissociation kinetics.

The peptide–lipid binding affinity of tachyplesin I, polyphemusin I, [I11A]tachyplesin I and [I11S]tachyplesin I was further examined with lipid bilayers prepared using lipid extracts from pathogenic and non-pathogenic bacteria, and from mammalian cells. The average lipid molar mass was determined from reported lipid profiles for each of the cell types (see Supplementary Table 1) and used to determine P/L at the end of peptide association, as above. Dose response curves (P/L) and representative SPR sensorgrams for 16 μM peptides are shown for EC958 (pathogenic bacteria), MG1655 (non-pathogenic bacteria) and BMMs (mammalian cells) in Fig. 5. Additional peptide–lipid binding curves are shown for *S.* Typhimurium, *B. humptydooensis* and RBC membranes in Supplementary Fig. 1.

**Fig. 5.**
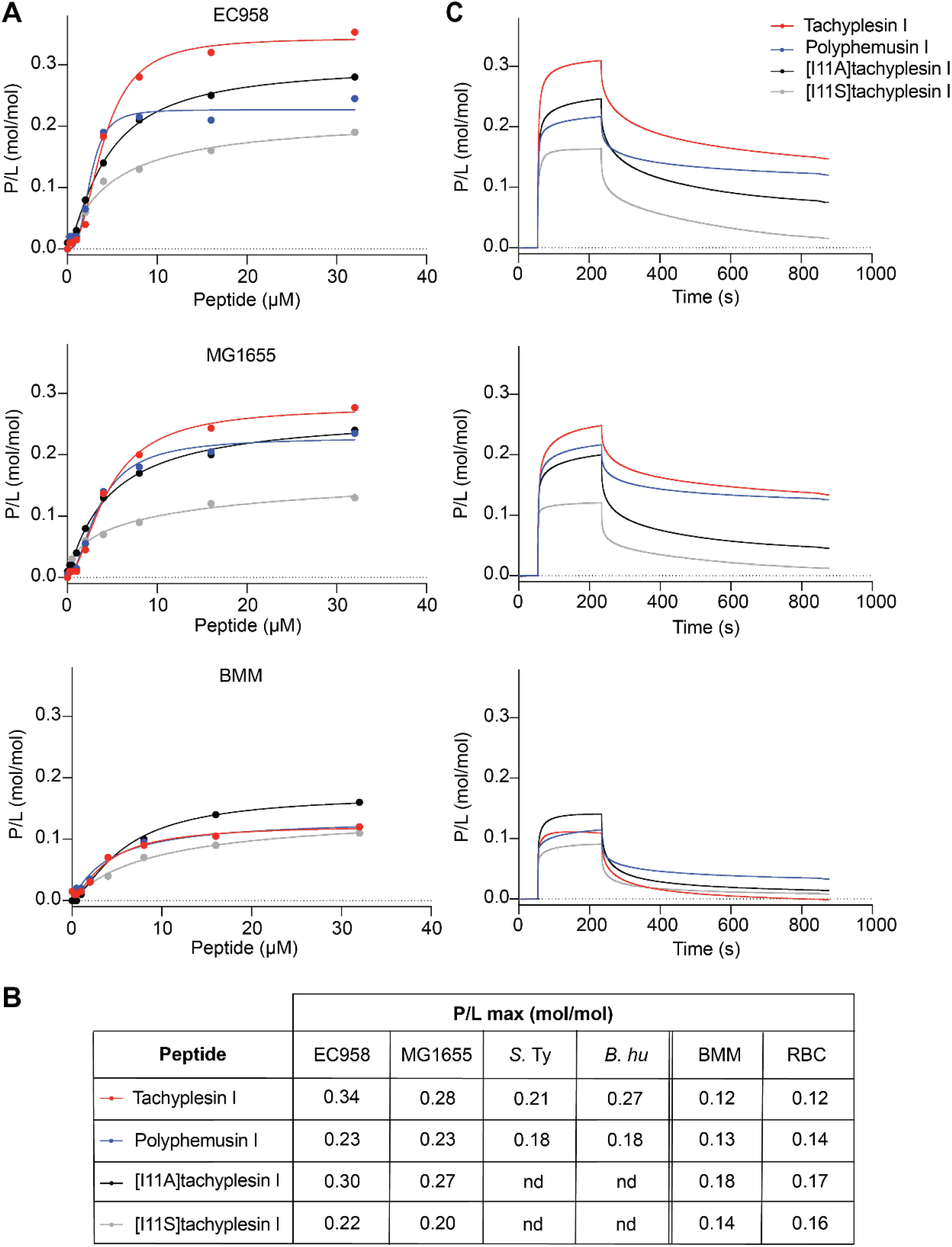
Membrane binding affinity and dose-response curves of tachyplesin I, polyphemusin I, [I11A]tachyplesin I and [I11S]tachyplesin I toward membrane extracts from bacterial and mammalian cells. Membranes extracts were deposited on an L1 chip. SPR sensorgrams were obtained for peptides injected over lipid bilayers for 180 s, with dissociation monitored for 600 s. The response units at the end of the association phase were converted to peptide-to-lipid ratios (P/L (mol/mol)). (A) Dose response curves allow comparison of peptide binding to EC958, MG1655 and BMMs membrane extracts. Peptide binding to *S*. Typhimurium (*S*. Ty), *B. humptydooensis* (*B. hu*) and RBC membranes are shown in Supplementary Figure 1. (B) The maximum P/L ratio (P/L max) was determined by fitting P/L doses response curves (saturation binding with Hill slope, GraphPad Prism 8), nd denotes no data. (C) Representative sensorgrams for 16 µM peptide show peptide-membrane binding characteristics.

All tested peptides bound to bilayers prepared using membrane lipids from bacterial cells with higher affinity than those prepared with lipids from BMMs or RBCs (Fig. 5B and Supplementary Fig. 1). Tachyplesin I had the highest maximal binding to bilayers composed with EC958 and MG1655 lipids, whereas [I11S]tachyplesin I had the lowest maximal binding to all of the tested bilayers with bacterial lipid membrane extracts (Fig. 5A and 5B). Binding to bilayers composed of other bacterial and mammalian membrane extracts (Supplementary Fig. 1) confirm greater affinity to lipid bilayers composed with bacterial lipids compared to mammalian cell lipids.

In general, the peptides showed higher binding affinity toward lipid bilayers with higher amounts of anionic phospholipids (including those with PG headgroups), as shown with bilayers composed with bacterial versus mammalian cell lipids, and with relative differences in the cytotoxic activity of peptides to corresponding cell types. Greater variability was observed between different peptides across the tested lipid bilayers composed with cell membrane extracts, than was observed for the model membranes composed with synthetic lipids, which is consistent with the higher degree of complexity present for the cell membrane extracts. However, the trend confirms that binding to anionic PG headgroups is likely to play an important role in antimicrobial activity of the peptides.

Differences in peptide–lipid binding affinity of the four peptides were less evident for model membranes (see P/L max Fig. 4B) and membrane extracts (Fig. 5B). Therefore, differences in antimicrobial activity or cell toxicity of tachyplesin I and analogues, and polyphemusin I (see Table 1) were not directly explained by peptide–lipid binding affinity. However, the > two-fold reduction in toxicity of [I11A]tachyplesin I and [I11S]tachyplesin I toward BMMs and RBCs might be explained by the smaller size of Ala, and increased polarity of Ser side chains compared to Ile at position 11, (opposite end of the peptide compared to membrane-inserting Trp at position 2) that reduce hydrophobic interactions as the peptide penetrates mammalian cell membranes.

### Membrane permeabilization

To determine whether the antimicrobial activity of the peptides involves bacterial membrane permeabilization, bacterial cells were treated with tachyplesin I, polyphemusin I, [I11A]tachyplesin I and [I11S]tachyplesin I, then exposed to SYTOX™ Green nucleic acid stain. SYTOX™ Green is impermeant to intact membranes, however, it readily penetrates cells with compromised membranes and binds to nucleic acids, causing >500-fold fluorescence enhancement. The percentage of bacteria cells that were permeabilized after treatment with peptide was determined by monitoring the fluorescence of SYTOX™ Green using flow cytometry. Dose response curves prepared using serial dilutions, starting from the MIC of peptide for each strain (Table 1), are shown in Fig. 6. Isopropanol was used as a control and shown to permeabilize 100% of bacterial cells.

**Fig. 6.**
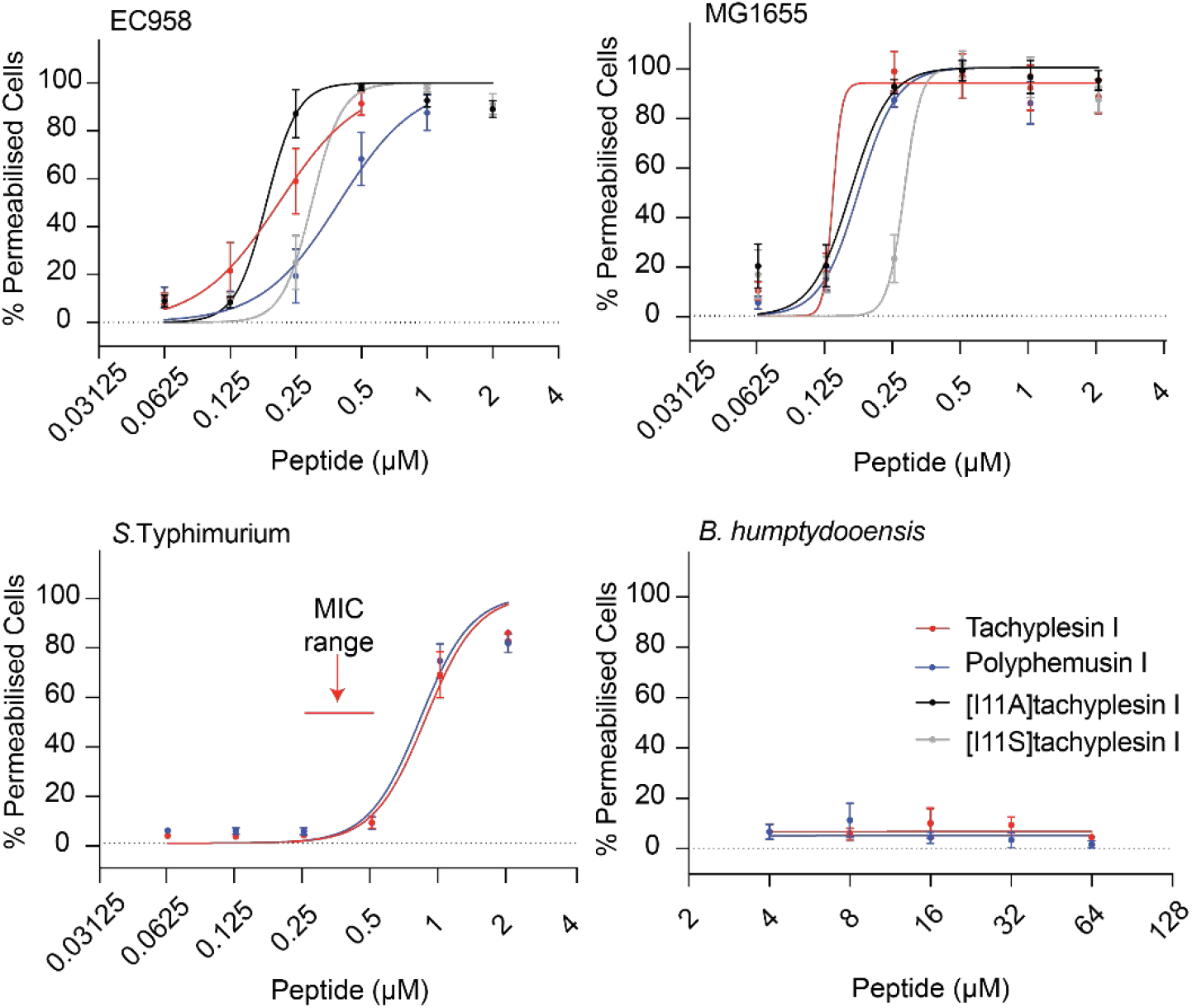
Comparison of peptide concentrations required to permeabilize bacterial cells. Bacterial cells were treated with serial dilutions of tachyplesin I, polyphemusin I, [I11A]tachyplesin I or [I11S]tachyplesin I and incubated in a shaking incubator for 1 h at 37°C. SYTOX® Green added to each sample only enters cells with damaged membranes to bind DNA. Fluorescent cells were detected using flow cytometry, excitation = 488 nm and emission 530/30 nm. 100,000 gated events were recorded for each condition. Dose response curves show the percentage of permeabilized cells, normalized using a control sample with cells treated with isopropanol (100% of permeabilized cells) and plotted using GraphPad Prism 8. Data represent the mean ± SEM from a minimum of three independent experiments. The red arrows on *S.* Typhimurium indicate the MIC.

EC958 and MG1655 cells were completely permeabilized by treatment with tachyplesin I, polyphemusin I, [I11A]tachyplesin I and [I11S]tachyplesin I at concentrations equal to the MICs (the highest concentrations tested for each peptide in this assay, Fig. 6). Concentrations below the MICs were found to cause permeabilization of a significant percentage of bacterial cells. For example, [I11A]tachyplesin I, which inhibits 100% of EC958 growth at 0.5–2 µM, permeabilizes approximately 100% of cells at 0.25 µM. [I11A]tachyplesin I permeabilized *E. coli* with the highest efficiency, while the lowest efficiency was observed for polyphemusin I. However, it is important to note that recorded cell populations only included intact cells.

In contrast to the high degree of permeabilization observed for *E. coli,* only ∼80% of *S.* Typhimurium cells were permeabilized following treatment with 1–2 µM polyphemusin I (equivalent to the MIC), and concentrations greater than the MIC for tachyplesin I. *B. humptydooensis* membranes appeared to remain intact, despite treatment with up to 64 µM tachyplesin I or polyphemusin I (Fig. 6).

### Peptide internalization of host cells

For peptides to be effective at targeting intracellular bacteria, they must be able to penetrate host cell membranes at concentrations that are toxic to the bacteria (e.g. EC958 0.5–2 μM, see Table 1), but are not toxic to host cells. The ability of tachyplesin I, polyphemusin I, [I11A]tachyplesin I and [I11S]tachyplesin I to internalize BMMs was examined using flow cytometry with peptides labelled with Alexa Fluor® 488, from a starting concentration of 1 µM. The percentage of fluorescent BMMs was determined in the absence and presence of trypan blue (TB, Fig. 7A), a membrane impermeable quencher of fluorescence. The identical fluorescence percentage in the absence and presence of trypan blue indicates that all the cells have their cell membrane intact, and all cells have internalized fluorescent peptide (and therefore not accessible to the quencher). All four peptides internalized into ∼100% BMMs at 1 μM (Fig. 7A). The [I11A] and [I11S]tachyplesin I analogues internalized more efficiently into BMMs, with > ten-fold lower concentrations required to internalize 50% of cells compared to tachyplesin I (Fig. 7A).

**Fig. 7.**
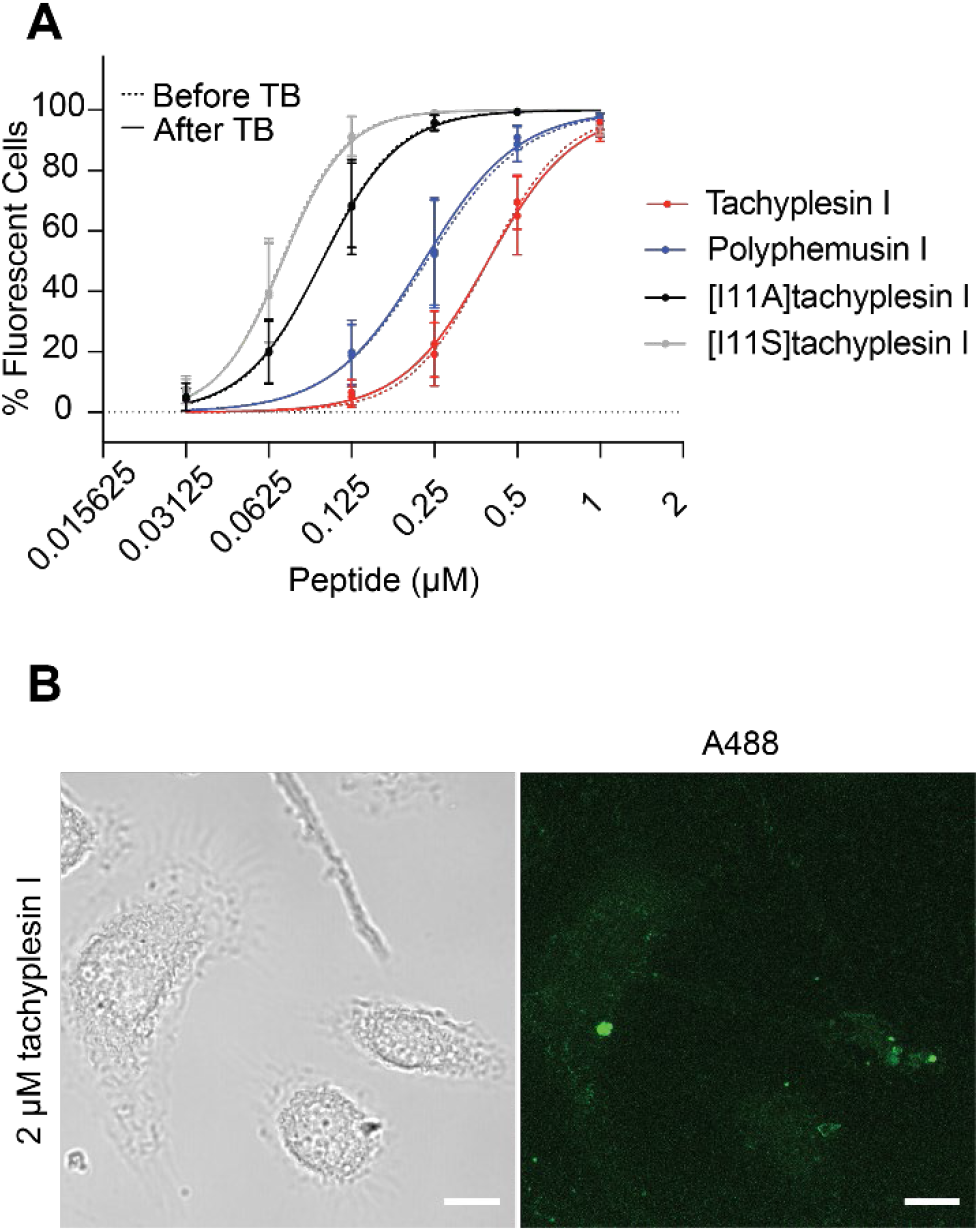
Internalisation of fluorescently labelled peptides into BMMs. (A) Tachyplesin I, polyphemusin I, [I11A]tachyplesin I and [I11S]tachyplesin I were labelled with Alexa Fluor ® 488 and incubated with BMMs for 1 h. Mean fluorescence emission intensity and the population of fluorescent cells were detected using flow cytometry with excitation at 488 nm and emission at 530/30nm. Trypan blue (TB) was added to quench the fluorescence of solvent accessible peptide (e.g. bound to the cell surface). The fluorescence emission intensity of 10,000 gated events were screened per peptide and the percentage of fluorescent cells were plotted, compared to unstained cells. Data represent the mean ± SEM from a minimum of three independent experiments. (B) BMM cells were plated onto micro Nunc™ Lab-Tek™ Chambered coverglass and treated with 2 µM tachyplesin I labelled with Alexa Fluor® 488 (green) and 100 µg/mL dextran-TMRE for 15 min. The medium was removed and replaced with FluoroBrite DMEM media. Cells were incubated at 37 °C and 5% CO2 during imaging. Confocal images were acquired using a Zeiss LSM 880 and are displayed here as brightfield and a maximum intensity projection. Images were processed and cropped in Fiji. Scale bar represents 10 µm.

Internalization of tachyplesin I was verified using confocal fluorescence microscopy of live BMMs (Fig. 7B). After treatment with 2 µM A488-tachyplesin I for 15 min, simultaneous uptake of peptide and dextran-TMRE was observed (see Supplementary Movie 1). The presence of fluorescent peptide and dextran-TMRE in discrete vesicles suggests that the predominant mechanism of cell entry under these conditions was via endocytosis. The more disperse fluorescence observed in some cells suggests that tachyplesin I can access the cytosol of BMMs within minutes. It is still not clear whether the cytosolic localization is a result of direct entry of peptide through the plasma membrane of BMMs, or whether the peptide within the vesicles is able to escape into the cytosol. It is possible that the observed cytosolic tachyplesin I is a result of both mechanisms of cell entry, similar to observations for other CPPs including the β-hairpin peptide cyclic gomesin [82] and cyclotide kalata B1 [67].

### Localization of tachyplesin I analogues inside BMMs infected with EC958

[I11A]tachyplesin I and [I11S]tachyplesin I, the two peptides that showed higher internalization efficiency (Fig 7A), were used to investigate if they could target intracellular bacteria. BMMs were infected with EC958 and treated for 1 h with peptides at the highest concentration that was nontoxic to BMMs (8 μM). EC958 were observed within intracellular vesicle compartments (Fig. 8, DAPI and TMRE panels) in agreement with our previous findings [26]. Treatment of infected BMMs with 8 µM labelled [I11A]tachyplesin I for 1 h resulted in a punctate appearance of fluorescence inside cells, suggesting that like tachyplesin I (Fig 7B), [I11A]tachyplesin I locates mainly within vesicles (Fig. 8, A488 panel and merge inset). In contrast, treatment of infected BMMs with 8 µM labelled [I11S]tachyplesin I for 1 h produced diffuse fluorescence throughout the BMM cytosol and strong co-localization with intracellular EC958 bacteria (Fig. 8).

**Fig. 8.**
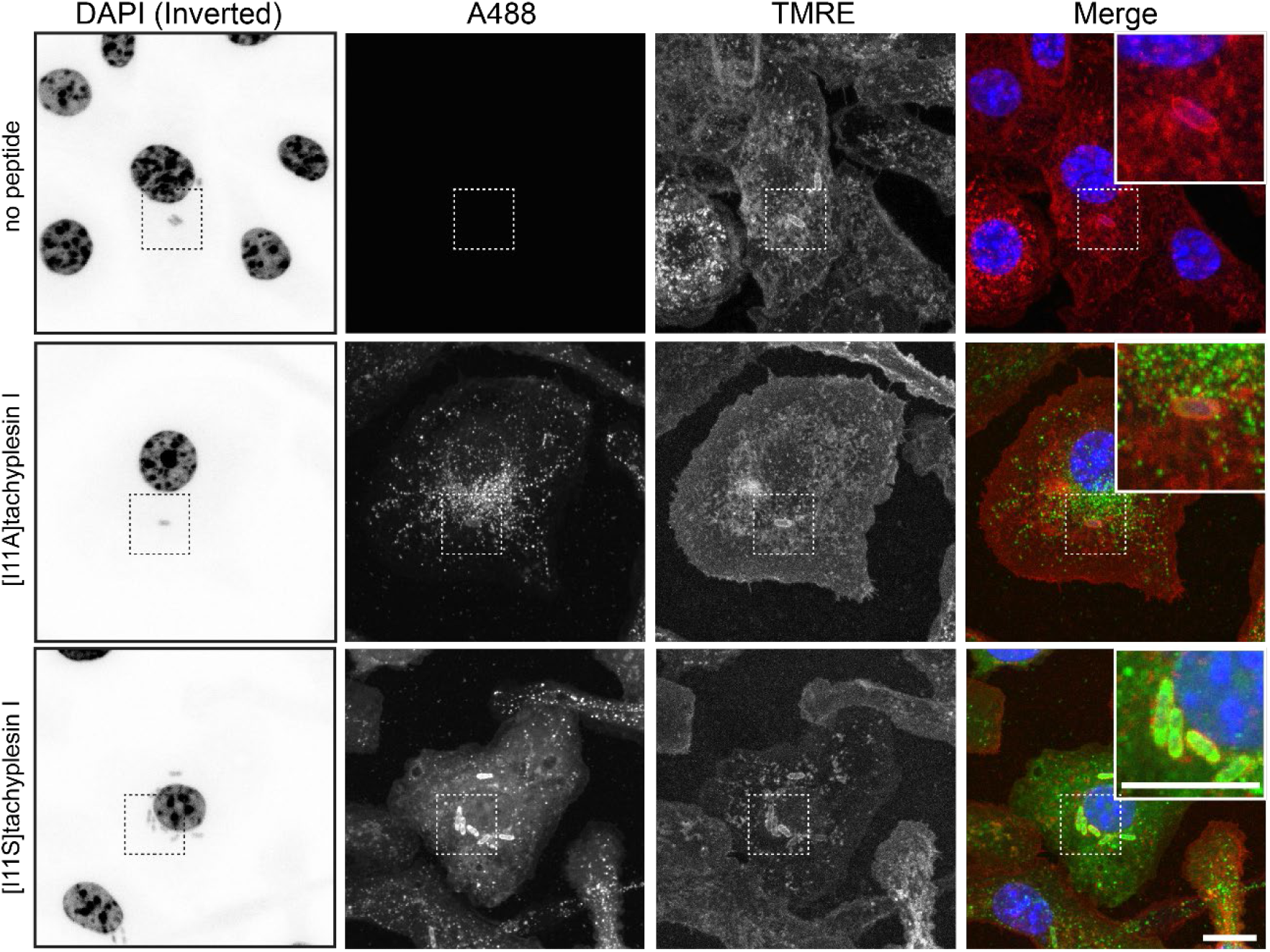
Intracellular localization of tachyplesin analogues in BMM cells infected with EC958. [I11A]tachyplesin and [I11S]tachyplesin were labelled with Alexa Fluor®488 (A488) (green). BMM cells were plated onto micro Nunc™ Lab-Tek™ Chambered coverglass and infected with EC958 (multiplicity of infection 1:100) for 2 h. Infected cells (media aspirated after treatment with 200 µg/mL gentamicin) were treated with 8 µM peptides for 1 h. The media was removed, and cells were fixed with 4% paraformaldehyde for 15 minutes. The fixative was removed and replaced with PBS and cells were stained with WGA-TMRE (staining the cell membranes, red) and DAPI (staining the DNA, blue). Untreated (no peptide) infected cells were included as a control. Confocal images were acquired using a Zeiss LSM 880 and are displayed here as maximum intensity projections. For clarity, the DAPI signal has been inverted. Images were processed and cropped in Fiji. Scale bar represents 10 µm.

### Investigating peptide ability to enhance zinc toxicity

Macrophages employ a variety of antimicrobial responses against pathogens. One mechanism relies on the modulation of metal ion concentrations, inducing starvation or direct toxicity, to eradicate bacteria during infection [49, 83–85]. In particular, zinc is essential for maintaining structure and catalytic activity of bacterial enzymes [86] and plays a role in signal transduction [87]. However, excessive zinc is cytotoxic, and immune cells can mobilize and deliver toxic concentrations of zinc to intracellular bacterial pathogens [47, 88].

Genome-wide analysis of EC958, employing a transposon directed insertion site (TraDIS) approach, identified a number of genes critical for survival in the presence of high zinc concentrations. Notably, this list included the bacterial zinc exporter ZntA, that counteracts intracellular zinc accumulation, along with other proteins required for maintaining cell membrane integrity [49]. Thus, due to the ability of tachyplesin I and polyphemusin I to bind and perturb bacterial membrane integrity, we hypothesized that co-treatment with zinc and peptides might have an additive antimicrobial effect, and that an EC958 strain lacking the zinc exporter ZntA (EC958Δ*zntA*) may be more sensitive than the wild type strain (EC958 WT) to zinc stress in presence of the peptides.

To test this hypothesis, a co-treatment analysis was performed with sublethal concentrations (below the MIC, see Table 1) of tachyplesin I, polyphemusin I, [I11A]tachyplesin I and [I11S]tachyplesin I and 0.1 mM zinc, which does not significantly inhibit EC958 WT or EC958Δ*zntA* growth (Fig. 9 and Supplementary Fig. 2). Except for a slight delay in onset of bacterial growth with polyphemusin I, treatment with a sublethal dose of peptide alone did not alter bacterial growth. Conversely, growth inhibition was observed after 6 h of co-treatment with 0.1 mM zinc and polyphemusin I, [I11A]tachyplesin I, or [I11S]tachyplesin I, which was more pronounced for EC958Δ*zntA* compared to EC958 WT (Fig. 9, 300 min). Despite the observed growth inhibition, co-treated EC958 WT returned to within 80–90% of the bacterial density that was observed for untreated bacteria after 12 h.

**Fig. 9.**
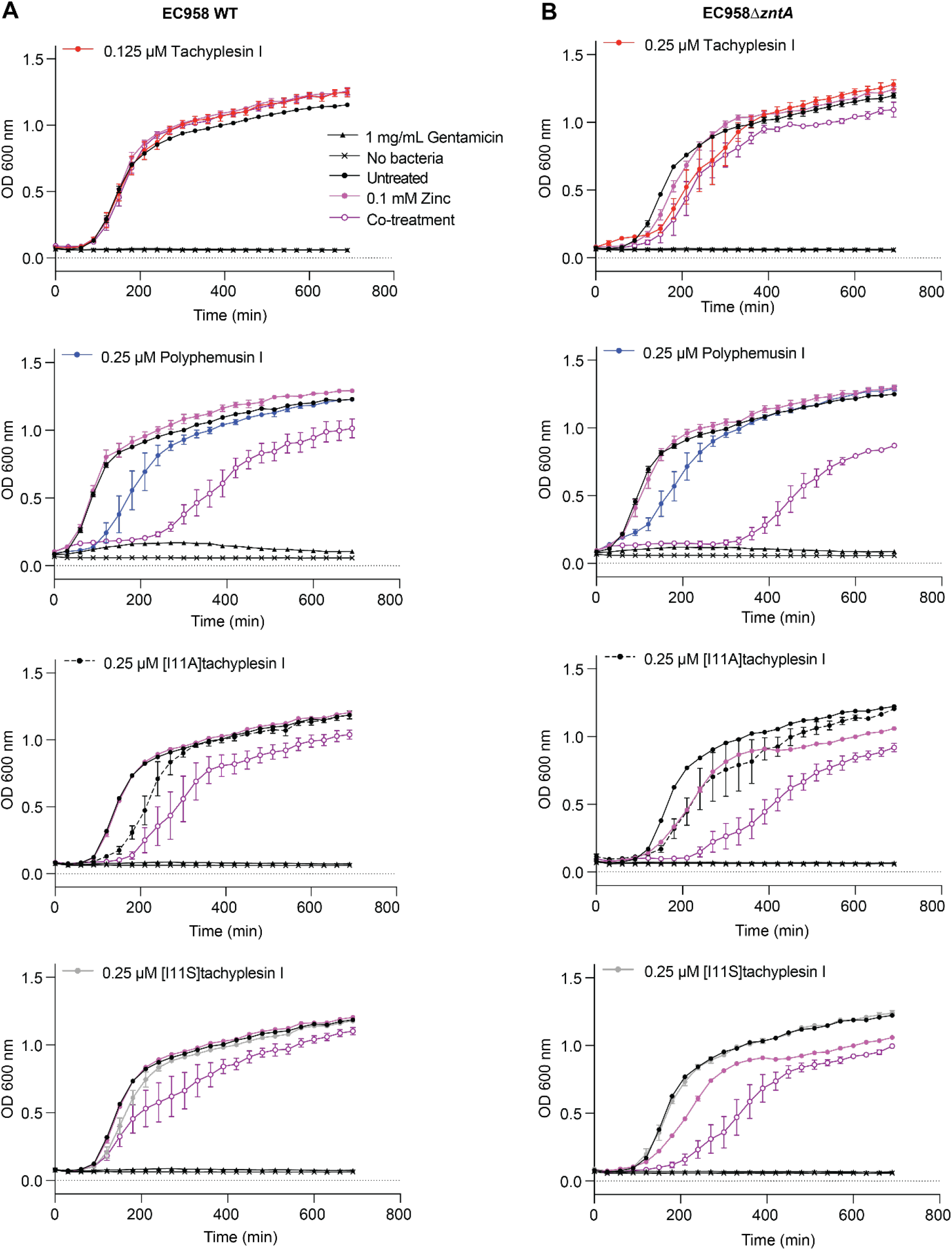
Treatment of EC958 WT and EC958Δ*zntA* with sublethal concentrations of peptides enhances zinc toxicity. Sublethal concentrations of tachyplesin I, polyphemusin I, [I11A]tachyplesin I or [I11S]tachyplesin I were determined from MIC data (Table 1). EC958 WT (A) and EC958Δ*zntA* (B) were treated with peptides in the presence and absence of 0.1 mM zinc. Samples were incubated at 37°C inside a PolarStar Omega plate reader and optical density was monitored by measuring absorbance at 600 nm every 30 min for 12 h. Controls included bacteria treated with 0.1 mM zinc, 1 mg/mL gentamicin (for 100% growth inhibition) or untreated. Data represent the mean ± SEM from a minimum of three independent experiments.

To investigate whether the observed additive effects were specific to zinc, EC958 WT was treated with 0.25 µM polyphemusin in the presence of copper or iron. Copper accumulation within the phagolysosome is thought to contribute to bacterial killing by mechanisms such as inhibition of bacterial metabolic processes, and by causing damage to proteins, lipids and DNA [83]. Iron is required by bacteria for survival as it participates in major biological functions, and it is important that it is maintained at non-toxic levels and in non-toxic forms [89, 90]. Indeed, accumulation of Fe(III) is reported to alter membrane integrity and lead to DNA damage via an excess production of hydroxyl radicals [91]. However, under similar experimental conditions to zinc, no additive effects were observed for polyphemusin I with either copper or iron at concentrations up to 0.5 mM (Supplementary Fig. 3). Together the peptide–metal ion co-treatment data suggest that polyphemusin I, [I11A]tachyplesin I and [I11S]tachyplesin I exert an additive effect on EC958 that is specific to zinc. Further mechanistic studies, including investigation into the effect of sublethal concentrations of peptide on expression levels of zinc exporter ZntA, and the effect on zinc stress reporter strains of EC958 [49] are warranted.

### The effect of peptides in an *in vitro* intracellular infection model

We established that tachyplesin I, polyphemusin I, [I11A]tachyplesin I, and [I11S]tachyplesin I exhibit antimicrobial activity toward EC958 WT and EC958Δ*zntA* at concentrations that are non-toxic to BMMs (Table 1); that the peptides can enter BMMs without damaging their membrane, at concentrations that permeabilize bacteria; and that sublethal concentrations of the latter three peptides enhance susceptibility of planktonic bacteria to zinc toxicity. EC958 can evade macrophage-mediated zinc toxicity, through mechanisms that may involve a higher resistance to zinc toxicity but also the capacity to protect themselves in LAMP1^+^ vacuoles [49]. Notably, EC958Δ*zntA*, which lacks its zinc exporter, can still subvert macrophage antibacterial responses and survive intracellularly. Therefore, EC958 WT and EC958Δ*zntA* were included in *in vitro* infection assays to determine whether peptide treatment enhances the zinc toxicity to affect intracellular bacterial survival.

To measure the effect on intramacrophage bacterial load, BMMs infected with EC958 WT or EC958Δ*zntA* were treated with peptide at concentrations that were lethal to bacteria (>MIC) but non-toxic to BMMs (< CC50, see Table 1). After 6 h and 12 h post-infection, BMMs were lysed and CFU were counted to determine the number of viable intracellular bacteria in all conditions. Treatment of BMMs with 0.5 µM tachyplesin I or 2 µM polyphemusin I did not reduce the number of intracellular EC958 WT at 6 h or 12 h post-infection (Fig 10A). However, when using the same concentration of polyphemusin I (2 µM) or slightly increased tachyplesin I (4 µM) on BMM infected with EC958Δ*zntA,* there was a trend for reduced bacterial loads after 12 h of infection, suggesting an enhancement of zinc toxicity (Fig. 10B). More dramatically, treatment with a higher concentration (8 µM) of the less toxic analogues, [I11A]tachyplesin I and [I11S]tachyplesin I, significantly reduced the bacterial load of both EC958 WT and EC958Δ*zntA* at both timepoints. Notably, the bacterial load of BMMs treated with [I11S]tachyplesin I was reduced to <5% of the control treatment after 10 h of peptide treatment (12 h post infection). We confirmed that this significant reduction in the number of bacteria (CFU) was not associated with toxicity to BMMs, as there was no increase in cell death measured by lactate dehydrogenase (LDH) released into the culture supernatant (Fig. 10C). The absence of toxicity to infected BMMs at these peptide concentrations is also consistent with uninfected BMM toxicity data (Table 1).

**Fig. 10.**
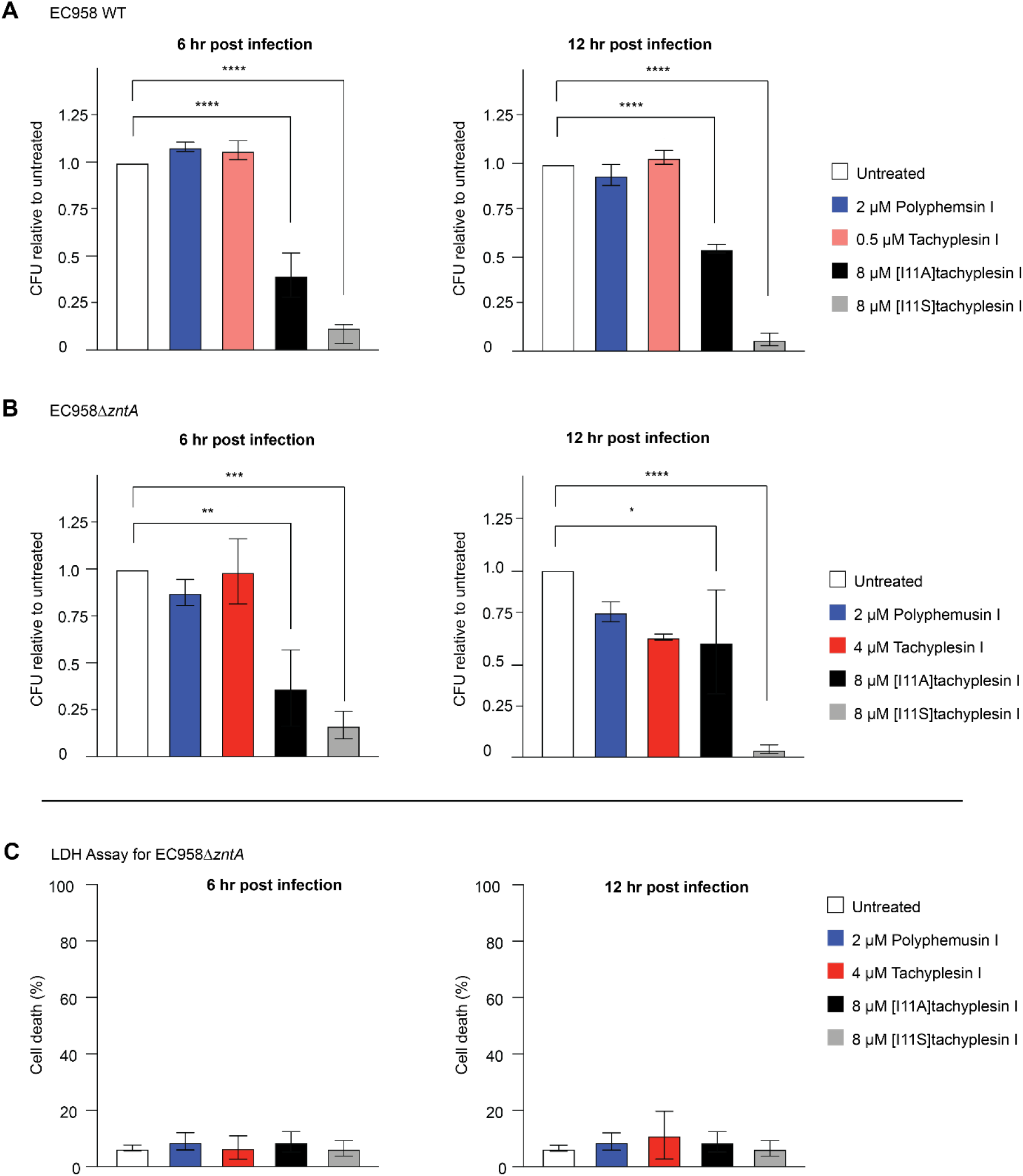
Reduction of intracellular bacterial load following treatment of BMMs with peptides. BMMs were infected with either EC958 WT or EC958Δ*zntA* for 1 h (multiplicity of infection (MOI) 1:100). Extracellular bacteria were removed by removing the supernatant and treating the replaced media with gentamicin (200 µg/mL final concentration) for 1 h. Media was removed and replaced with media containing 20 µg/mL gentamicin and peptides at MICs (tachyplesin I and polyphemusin I) or the highest possible non-toxic dose of 8 µM ([I11A]tachyplesin I and [I11S]tachyplesin I). After 4 h or 10 h of peptide treatment (equivalent to 6 h or 12 h post infection), BMMs were lysed with 0.01% (*v/v*) Triton X-100. Supernatant was diluted and plated on agar. Agar plates were incubated at 37°C for 12 h. Colonies were counted and converted to CFUs. The percentage of CFU compared to untreated controls were plotted ± SEM from three independent experiments for (A) EC958 WT or (B) EC958Δ*zntA*. (C) A lactate dehydrogenase activity assay (LDH) was performed in parallel to the infection assay. BMMs were infected with MOI 1:100 EC958 WT or EC958Δ*zntA* and treated in the same manner as the infection assay. The LDH assay was performed as per manufacturer’s instructions, taking samples at 6 h and 12 h post infection. Absorbance at 480 nm was read on a Tecan infinite M1000Pro and the percentage cell death was calculated using the ratio of media subtracted conditions divided by the subtracted absorbance of 100% lysis controls. The LDH assay for EC958Δ*zntA* is presented here. Significance for all panels was determined by one-way ANOVA. P values are denoted by: * P ≤ 0.05, ** P ≤ 0.01, **** P ≤ 0.0001.

Taken together with the planktonic co-treatment study, it is proposed that the enhanced intracellular antimicrobial activity of these peptides against EC958Δ*zntA* compared to EC958 WT may be due to the additive effect of intramacrophage zinc toxicity, and the diminished ability of EC958Δ*zntA* to overcome this effect. This finding is important as it was previously shown that UPEC evades the zinc toxicity response within human macrophages [49]. Species differences could be one explanation, as human and murine macrophages trigger evolutionarily divergent inflammatory responses [92], but a more exciting possibility is that the low level zinc exposure of UPEC in macrophages, usually non-toxic, is amplified when CPPs exert their effect on bacterial membranes.

### Conclusions

The growing problem of antibiotic resistance is made more challenging by the intracellular lifestyle of some pathogenic bacteria. To effectively control this subset of bacteria, new therapies must be able to reach and kill bacteria that sequester within vesicles to survive and replicate, at concentrations that do not damage host cells. Here we demonstrate the potential for host defence peptides, especially β-hairpin peptides from horseshoe crabs, tachyplesin I and polyphemusin I, to inhibit the growth of UPEC bacteria in both extracellular and intracellular environments. Moreover, a single amino acid substitution to produce [I11S]tachyplesin I reduced toxicity towards host BMM cells and greatly increased the ability to access intracellular bacteria to clear infection with multidrug resistant UPEC strain EC958.

Polyphemusin I, tachyplesin I and analogues [I11A]tachyplesin I and [I11S]tachyplesin I inhibit bacterial growth at concentrations at least ten-fold lower than concentrations that are toxic to mammalian cells. This selectivity is related to their preferential binding to biological membranes that are rich in negatively charged phospholipid headgroups. Furthermore, these peptides permeabilize membranes of both pathogenic and non-pathogenic *E. coli* at sublethal concentrations, to inhibit bacterial growth. The ability of these peptides to permeabilize membranes at sublethal concentrations is particularly important as EC958 is known to avoid antimicrobial pathways and to remain quiescent in macrophages for long periods of time. Indeed, compared to the non-pathogenic *E. coli*, EC958 manifests a high tolerance to zinc toxicity, through both evasion and resistance. Our data suggest that the CPPs polyphemusin I, [I11A]tachyplesin I and [I11S]tachyplesin I, target and fragilize bacterial membranes, enabling sensitivity to zinc at concentrations that are usually non-toxic for EC958.

[I11A]tachyplesin I and [I11S]tachyplesin I analogues provided at least a two-fold increased therapeutic window compared to native tachyplesin I, thereby affording treatment of BMMs infected with UPEC at higher concentrations, and subsequently significant reduction of intramacrophage bacterial load. The basis of this improved efficacy remains unclear, as overall structure and affinity of the peptide analogues for model and extracted bacterial membranes was similar to that of native tachyplesin I and polyphemusin I. Nevertheless, these data highlight the potential for host defence peptides as new modalities for targeting bacteria with intracellular niches. Overall, this study contributes to the growing body of evidence that antimicrobial peptides are an attractive alternative to conventional small molecule drugs, especially for pathogenic bacteria with an intracellular lifestyle.

## DECLARATIONS

### Author Contributions

Anna Amiss, Nicole Lawrence, Sónia Troeira Henriques, and Ronan Kapetanovic contributed to the study conception and design. Material preparation, data collection and analysis were performed by Anna Amiss, Jessica Von Pein, Jessica Webb, Nicole Lawrence, Nicholas Condon, and Peta Harvey. Funding and resources were provided by Nicole Lawrence, Sónia Troeira Henriques, David Craik, Bart Currie, Matthew Sweet, and Mark Schembri. The manuscript was written by Anna Amiss, Nicole Lawrence, Sónia Troeira Henriques, and Ronan Kapetanovic. All authors commented on the previous versions of the manuscript and read and approved the final manuscript. The authors would like to thank Dr Kaustav Gupta at The Institute for Molecular Bioscience, Centre for Inflammation and Disease Research and Australian Infectious Diseases Research Centre, The University of Queensland, for harvesting and culturing bone marrow macrophages.

### Availability of data and material

Peptide sequences are provided herein. Average data are provided, with processing and curve fits as reported for specific experiments.

### Ethics approval

The University of Queensland Institutional animal ethics committee approved the use of primary mouse cells, herein referred to as BMM cells (IMB/123/18). RBCs were collected from healthy adult donors following protocols approved by the Human Research Ethics Committees (University of Queensland approval number 2013000582).

