## Supplementary figures for "Modified defence peptides from horseshoe crab target and kill bacteria inside host cells"

| | | |
|---|---|---|
| Ronan Kapetanovic | | |
| Sónia Troeira Henriques | | Tel: +61 7 3443 7342 |
| Nicole Lawrence | | Tel: +61 7 3346 2020 |

### Supplementary Table 1: Average lipid mass for P/L calculations for lipid bilayers of cell membrane extracts

| | Cell membrane composition (%) [a] | | | | |
|---|---|---|---|---|---|
| **Lipid subtype** [b] | ***E.coli*** | ***S.* Typhimurium** | ***Burkholderia*** | **BMM** | **RBC** |
| Phospholalanine | - | - | - | - | 1.1 |
| Phosphocholine | - | - | - | 30.1 | 29.3 |
| Phosphoethanolamine | 77 | 92 | 64.9 | 21.6 | 27.6 |
| Phosphoglycerol | 9 | 5 | 12 % | - | - |
| Phosphoinositol | - | - | - | 6 | 0.6 |
| Phosphoserine | - | - | - | 6 | 14.9 |
| Cardiolipin | 14 | 3 | 9.9 | - | - |
| Sphingomyelin | - | - | - | 15.4 | 25.5 |
| Other | - | - | 13.2 | 20.9 | 1 |
| **Average mw (Da)** | **831.1** | **743.5** | **706.1** | **752.2** | **734.9** |

[a] Proportional representation of lipid subtypes was determined from previously reported cell membrane compositions for *Escherichia coli* (*E. coli*, represented by K-12) [1], *Salmonella enterica* serovar Typhimurium (*S.*Typhimurium) [2], *Burkholderia* (represented by *B. cenocepacia*) [3], murine bone marrow-derived macrophages (BMM)[4], and human red blood cells (RBC) [5].

[b] Molecular weight of representative lipids was used for determining the average molecular weight (mw) of the cell membrane extracts. 1-palmitoyl-2-oleoyl-sn-glycero-3 (PO)-phosphate (POPA, 696.5), PO-phosphocholine (POPC, 696.5 Da), PO-phosphoethanolamine (POPE, 717.5), PO-phosphoglycerol (POPG, 770.5), PO-inositol (POPI, 856.3), PO-phosphoserine (POPS, 783.5), Cardiolipin (1494.3), sphingomyelin (18:0 SM, 730.6), and 'other' components not used in the calculation.

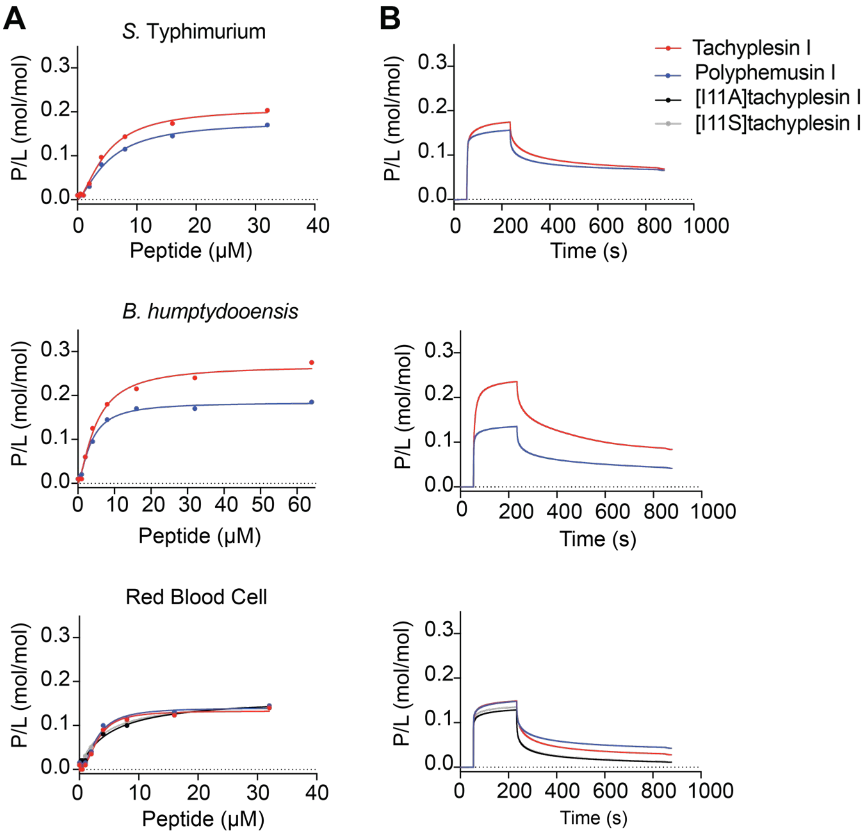


**Supplementary Fig. 1 Membrane binding affinity and dose-response curves of tachyplesin I, polyphemusin I, [I11A] tachyplesin I and [I11S] tachyplesin I toward membrane extracts**. Membranes from *S. Typhimurium*, *B. humptydooensis* and red blood cells were extracted and used to create a lipid bilayer on an L1 chip. SPR sensorgrams were obtained for peptides injected over lipid bilayers for 180 s, with dissociation monitored for 600 s. The response units at the end of the association phase were converted to peptide-to-lipid ratios (P/L (mol/mol)). (A) Dose response curves allow comparison of peptide binding to the membrane extracts. The maximum P/L ratio (P/L max) was determined by fitting P/L doses response curves (one site – specific binding with Hill slope, GraphPad Prism 8) and is shown in Figure 5B. (B) Representative sensorgrams for 16 µM peptide show peptide-membrane binding characteristics.

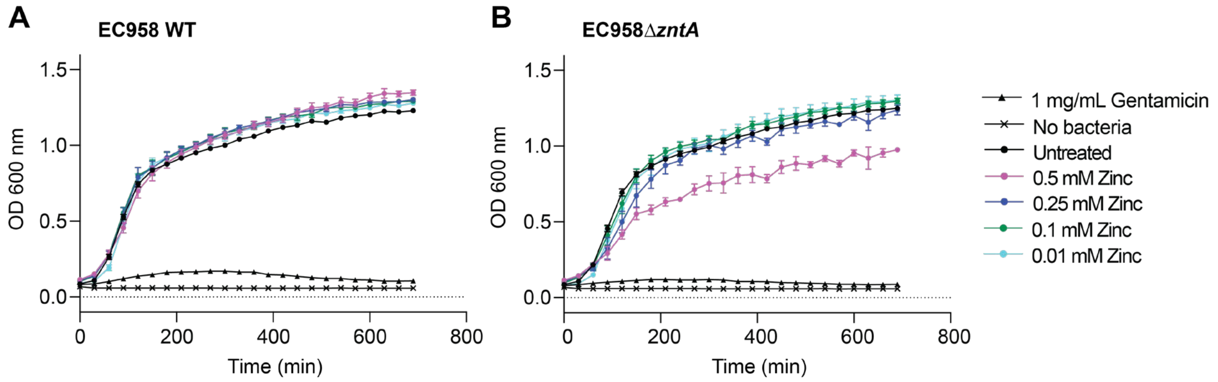


**Supplementary Fig. 2 The effect of zinc on the growth on EC958.** EC958 WT (A) and EC958Δ*zntA* (B) were treated with a range of of zinc sulfate concentrations. Samples were incubated at 37°C and absorbance readings at 600 nm were measured with a PolarStar Omega. Readings were taken every 30 min for a total of 12 h. Data represent the mean ± SEM from a minimum of three independent experiments.

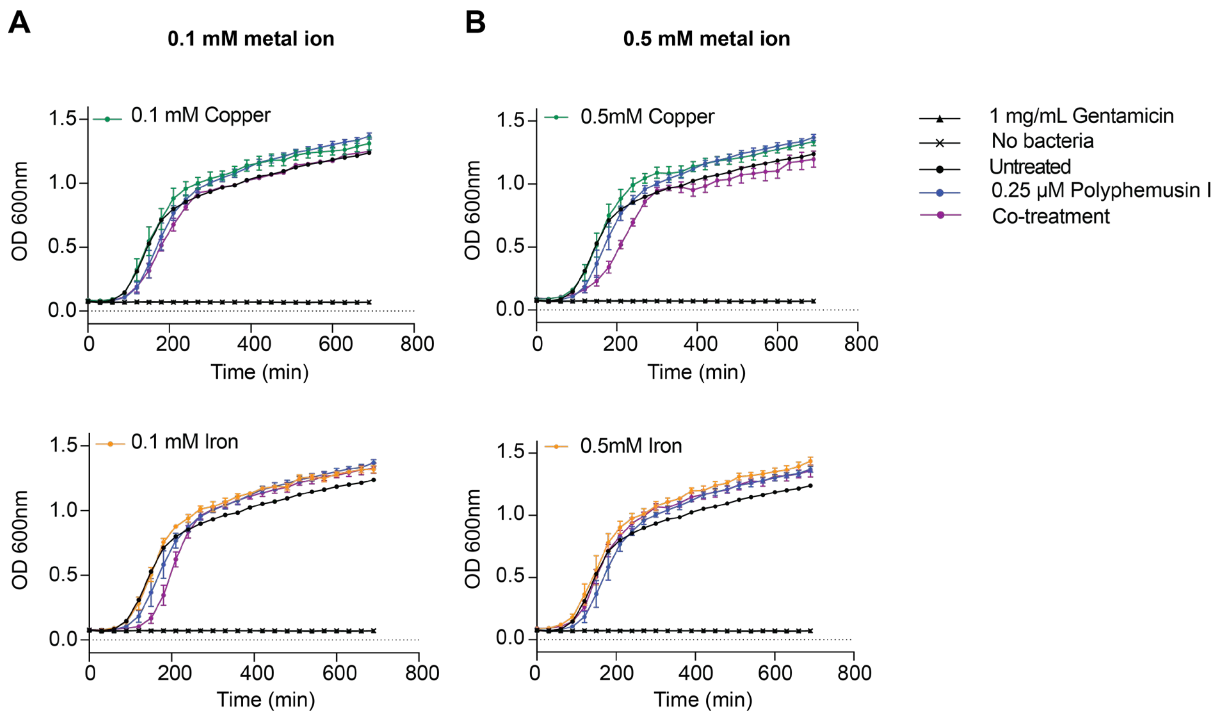


**Supplementary Fig. 3 Co-treatment of EC958 WT with 0.25 µM polyphemusin I with copper or iron.** Using a sublethal concentration of polyphemusin I against EC958 WT (from MIC determined in the peptide susceptibility screen, see Table 1). (A) EC958 WT cells were treated with 0.25 µM polyphemusin I ± 0.1 mM of copper sulfate or iron sulfate. (B) EC958 WT cells were treated with 0.25 µM polyphemusin I ± 0.5 mM of copper sulfate or iron sulfate Samples were incubated at 37°C and absorbance readings at 600 nm were measured with a PolarStar Omega. Samples were shaken prior to absorbance readings at 600 nm. Readings were taken every 30 min for a total of 12 hrs. Data was plotted using GraphPad Prism 8.0 and represents the mean ± SEM from a minimum of three independent experiments.
